# Subcellular dynamics studies reveal how tissue-specific distribution patterns of iron are established in developing wheat grains

**DOI:** 10.1101/2021.02.14.431124

**Authors:** Sadia Sheraz, Yongfang Wan, Eudri Venter, Shailender K Verma, Qing Xiong, Joshua Waites, James M Connorton, Peter R Shewry, Katie L Moore, Janneke Balk

**Affiliations:** School of Materials and Photon Science Institute, University of Manchester, Manchester, M13 9PL, UK; Department of Plant Sciences, Rothamsted Research, Harpenden, AL5 2JQ, UK; Bioimaging facility, Department of Computational and Analytical Sciences, Rothamsted Research, Harpenden, AL5 2JQ, UK; Department of Biological Chemistry, John Innes Centre, Norwich NR4 7UH, UK; School of Biological Sciences, University of East Anglia, Norwich NR4 7TJ, UK

**Keywords:** iron, pulse-chase, trafficking, nicotianamine, aleurone, wheat, NanoSIMS

## Abstract

Understanding iron trafficking in plants is key to enhancing the nutritional quality of crops. Due to the difficulty of imaging iron in transit, little is known about iron translocation and distribution in developing seeds. A novel approach, combining ^57^Fe isotope labelling and NanoSIMS, was used to visualize iron translocation dynamics at the subcellular level in wheat grain, *Triticum aestivum* L. We were able to track the main route of iron from maternal tissues to the embryo through different cell types. Further evidence for this route was provided by genetically diverting iron into storage vacuoles, as confirmed by histological staining and TEM-EDS. Virtually all iron was found in intracellular bodies, indicating symplastic rather than apoplastic transport. Aleurone cells contained a new type of iron body, highly enriched in ^57^Fe, and most likely represents iron-nicotianamine being delivered to phytate globoids. Correlation with tissue-specific gene expression provides an updated model of iron homeostasis in cereal grains with relevance for future biofortification efforts.

## Introduction

As widely consumed staple crops, cereals are important sources of mineral micronutrients, including iron and zinc which are essential for human health. However, deficiencies in these minerals affect large parts of the global population (WHO, 2013, 2015). The daily requirements for bioavailable iron and zinc are often not met in cereal-based diets for two reasons. First, iron and zinc are unevenly distributed in the grain, accumulating at high concentrations in the embryo (germ) and outer layers (bran), which are removed during polishing or milling. By contrast, both minerals are low in the starchy endosperm which constitutes about 70% of the grain and is preferentially consumed in human diets (as white wheat flour and polished rice, for example). Second, most of the iron and zinc in the embryo and aleurone layer is bound to phytic acid, forming insoluble complexes that have poor bioavailability. Attempts to biofortify cereal grains are focussing on increasing the total amounts of iron and zinc, changing their distribution and reducing phytic acid levels (Vasconcelos et al., 2017; Cominelli et al., 2020).

The molecular pathways of iron and zinc uptake from the soil into plant roots are relatively well understood, but we know little about mineral loading into the seeds and subsequent distribution to different tissues (Mari et al., 2020). Recent isotope labelling studies in Arabidopsis suggested that virtually all the iron in seeds was remobilized from senescing leaves (and other organs) with only an indirect contribution from uptake by the roots (Pottier et al., 2019). Studies using isotope pulse labelling have not yet been conducted in wheat, but a time course of mineral partitioning indicated that 77% of the iron in mature grain is remobilized from the shoot (Garnett and Graham, 2005), and that this process is regulated by NAC transcription factors (Uauy et al., 2006; Waters et al., 2009; Borrill et al., 2019).

Physical separation between the tissues of the mother plant and the seed (the filial generation) means that iron is secreted and then taken up again by the developing zygote (Mari et al., 2020). Nutrients are delivered to the developing grain via the vascular bundle which runs through the ventral crease in wheat, and the nucellar projection. The latter is all that remains from the nucellus which once surrounded the embryo sac, and comprises a dense group of transfer cells along the top of the vascular bundle (see Bechtel et al. (2009) for a description of wheat grain development). In the early stages of grain development, nutrients are secreted from the maternal transfer cells into a cavity that will fill up as the endosperm of the developing seed expands. The rapidly dividing endosperm cell mass differentiates into specialized cell types (Olsen, 2020), most notably a single outer layer of aleurone cells which differ from the other endosperm cells in lacking starch and accumulating protein, lipids, minerals and phytic acid. Periclinal divisions of the aleurone cells continue to form several layers of subaleurone cells which accumulate protein but not minerals. The aleurone cells that are in contact with the nucellar transfer cells in the tip of the crease differentiate to have a special function in nutrient transport. For this reason they are often called endosperm transfer cells, but to avoid confusion with the nucellar transfer cells we will use the term modified aleurone here, which is also commonly used in the literature (Evers, 1970; Borg et al., 2009).

The distribution of iron in biological materials has been visualised using histological staining, X-ray fluorescence (XRF), micro-proton induced X-ray emission (μ-PIXE) and laser-ablation inductively coupled plasma mass spectrometry (LA-ICP-MS). When applied to cereal grains, these techniques (>10 μm resolution) showed accumulation of iron in the crease, the aleurone layer and the scutellum of the embryo in wheat (Neal et al., 2013; Singh et al., 2013; De Brier et al., 2016) and a similar pattern in other cereals (Iwai et al., 2012; Takahashi et al., 2009; Detterbeck et al., 2020). Nanoscale Secondary Ion Mass Spectrometry (NanoSIMS) has also become a significant tool to visualise minerals at the subcellular level due to its unique capabilities of high spatial resolution (50 nm), high sensitivity (ppm and ppb for some elements) and detection of trace elements and isotopes (for example see Malherbe et al., 2016; Kopittke et al., 2020). During NanoSIMS analysis, the sample surface is impacted with a high-energy primary ion beam which causes sputtering of the surface and ejection of atoms and small molecules. Some of this sputtered material becomes ionised, referred to as ‘secondary ions’, which are detected and analysed by mass in a double focussing mass spectrometer. The instrument has two primary ion sources to generate either a caesium ion beam (Cs^+^), used for analysis of negative secondary ions or an oxygen ion beam (O^−^) to analyse positive secondary ions. Due to the design of the NanoSIMS, the secondary ions must have an opposite polarity to the primary ions. Up to 7 secondary ions can be detected simultaneously but these must be selected before acquiring the data (Hoppe et al., 2013). As the NanoSIMS operates under ultra-high vacuum, careful sample preparation is required to avoid redistribution of elements from their *in vivo* location (Grovenor et al., 2006).

Very few genes have been characterized to date that influence the distribution pattern of iron in seeds. One of those is the vacuolar iron transporter VIT. Distruption of this gene in Arabidopsis resulted in relocation of iron from provascular strands to the abaxial (lower) epidermis in the embryo which occupies most of the seed volume (Kim et al., 2006). In rice, mutation of either the *VIT1* or *VIT2* paralogue leads to iron accumulation in the embryo and depletion in the large endosperm of the cereal grain (Zhang et al., 2012; Bashir et al., 2013; Che et al., 2021) Conversely, overexpression of *TaVIT2* in the starchy endosperm of wheat grain leads to iron accumulation in this tissue and >2-fold more iron in white flour (Connorton et al., 2017).

Many questions remain regarding the mechanisms that determine how iron is translocated from the maternal vascular bundle into the developing seed and how this element is distributed within the seed as tissues differentiate and expand. To address these questions, we combined iron isotope labelling with NanoSIMS to compare the dynamics of iron distribution in developing wheat grains between a control and a *TaVIT2* overexpressing line. Our results revealed that the major route of iron is from the nucellar transfer cells through a zone of endosperm cells between the crease and the embryo, but that this pathway is disrupted by overexpression of *TaVIT2*. Different cell types displayed specific patterns of isotope-enriched vesicles and globoids, highlighting the different roles of each cell type in iron translocation.

## RESULTS

### Genetic characterization of *HMW-TaVIT2* line 22-19 for further study

Previously we generated 25 independent *Agrobacterium*-transformed wheat lines with one or more copies of the *HMW-TaVIT2* transgene, placing *TaVIT2* under the control of the starchy endosperm-specific promoter of the gene *High Molecular Weight subunit of glutenin 1D×5* (Connorton et al., 2017). Transcript levels of *TaVIT2* in developing grain correlated well with the number of transgene copies (factor 0.2, R^2^ = 0.597, p < 0.01). By contrast, the iron concentration in hand-milled flour was variable between lines with similar transgene copy numbers. To select one line for detailed study, we chose line 22-19 based on a representative pattern of iron staining in hand-dissected grain and a consistent >2-fold increase in the iron concentration in the white flour fraction. T-DNA copy number analysis of the T1 generation indicated that line 22-19 had more than one insert of tandemly arranged T-DNAs. Further selection gave a stable pattern of 8 – 10 T-DNA copies in the T2, suggesting the line was homozygous for a tandem array of 4 copies and possibly heterozygous for another single copy insertion (Suppl. Table S1).

To determine the position of the T-DNA insertion(s), TAIL-PCR (thermal asymmetric interlaced PCR) was performed with nested primer sets for either the left border (LB) or right border (RB) T-DNA sequence, in combination with an arbitrary degenerate (AD) primer. A 1.1-kb PCR product generated with the LB primer set and the AD3 primer was found consistently in transgenic plants but not in the non-transgenic control. Part of the sequence of the PCR product matched the LB and the other part matched the 5’UTR of *TraesCS4D02G046700* (Suppl. Fig. S1A, B). No specific PCR products were generated with the RB primer set with any of the 4 AD primers. The position of the T-DNA was verified with primers spanning the insertion site and the LB3 primer (Suppl. Fig. S1C). Analysis of DNA from 12 T1 siblings showed that the selected line (22-19-4) was homozygous for this T-DNA insertion, in agreement with the segregation pattern of T-DNA copies in the T2 generation.

*TraesCS4D02G046700* encodes a Ubc enzyme variant (Uev), belonging to a small gene family conserved in all eukaryotes. The closest rice homologue is *OsUEV1A (Os03g0712300)* which has 85% amino acid identity (Wang et al., 2017). Transcript levels of the wheat *UEV1A-4D* gene are slightly increased in developing grain, but this was also the case for the 4A and 4B homeologs (Suppl. Fig. S1D). Moreover, the wheat UEV1A-4A and - 4D homeologs are identical in amino acid sequence and have highly similar expression patterns (expVIP database, wheat-expression.com). Thus, the T-DNA insertion in line 22-19 does not appear to disrupt the expression of an essential gene.

### In TaVIT2 grain, the amount of iron is increased in a specific region of the endosperm and decreased in the embryo and aleurone cells

To further investigate differences in iron distribution as a consequence of overexpressing *TaVIT2* in the endosperm, T3 grains from line 22-19-4-5 (henceforth called TaVIT2) and from control plants were cut longitudinally and at two transverse positions before staining with Perls’ reagent (Fig. 1). Similar to other *TaVIT2* transformation events (Connorton et al., 2017), dense iron staining was found in the starchy endosperm surrounding the crease. Iron staining in this part of the endosperm was more intense in the proximal region close to the embryo than in the distal region (Fig. 1A, compare transverse section a and b), in a zone of cells equivalent to the Endosperm Adjacent to Scutellum (EAS) in maize kernels (Doll et al., 2020). By contrast, both the embryo and aleurone layer in TaVIT2 grain displayed a lower intensity of iron staining than in control grain. The difference in iron staining of the aleurone was also observed in higher magnification images using diamine benzidine (DAB)-enhanced Perls’ staining (Fig. 1B).

**Figure 1.**
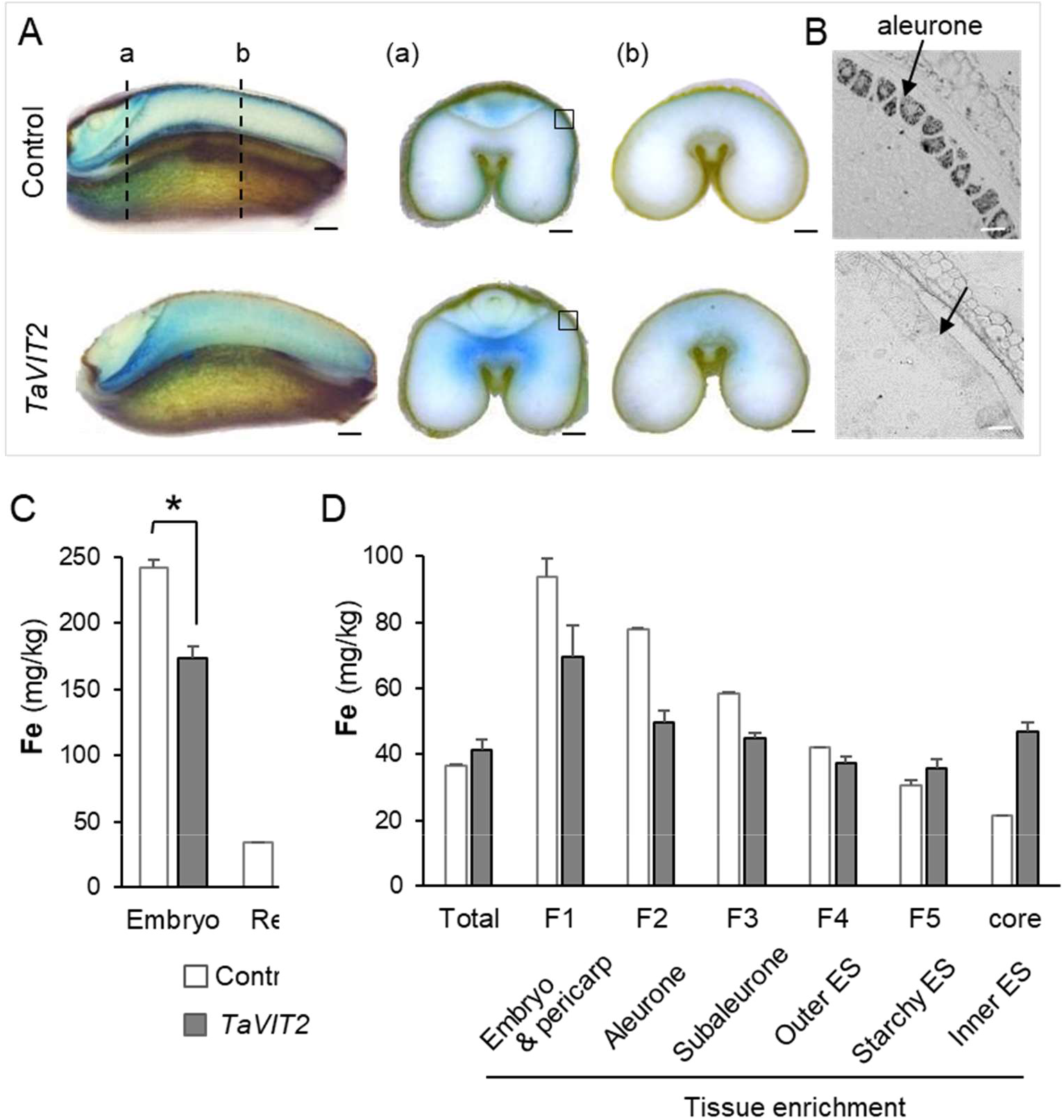
Distribution of iron in developing wheat grains overexpressing *TaVIT2* in the endosperm and non-transformed control. **A**. Grains were harvested 24 days after anthesis, hand-sectioned and stained for iron (blue) using the Perls’ staining method. The dashed lines (a, b) in the longitudinal sections on the left indicate the positions of the transverse sections on the right. Scale bars 0.5 mm. **B**. Thin sections of embedded material were stained with enhanced Perls’-staining. The grey-scale images show part of the aleurone cell layer marked by the square in A (a). Scale bars are 50 μm. **C**. Iron concentration per unit dry weight determined by ICP-MS. Embryos were hand-dissected from grains at 22 – 25 dpa. Values represent the mean ± SE of 3 biological replicates, which are pools of 40-50 embryos from one ear. *P < 0.05, Student t-test. **D**. Fractions obtained by pearling of mature grain were analysed for iron concentration (shown here) and other elements (Suppl. Fig. 3). Values represent the mean of 2 biological replicates of 30 g grain. Error bars represent half the difference between the measurements. The tissue enrichment in each fraction is based on (Tosi et al., 2018).

To quantify the decrease in iron content in the embryos, developing grains at 24 dpa were hand-dissected into two parts, embryo and the remaining tissues (rest), and the concentration of iron was measured by inductively coupled plasma mass spectrometry (ICP-MS). Iron was significantly decreased, by 28.5%, in embryos of TaVIT2 grains compared to control embryos, with a concentration of 173 ± 20 mg kg^−1^ dry weight compared to 242 ± 9 mg kg^−1^ in control (p > 0.03, see Fig. 1C). However, germination tests of TaVIT2 grain showed no effect on germination or on early seedling growth in alkaline soil (Suppl. Fig. S2).

The iron contents of the aleurone layer and endosperm of mature grains were estimated by analysis of pearling fractions. This technique uses abrasion to remove material from the outer layers to the inner core of the grain. Sequential cycles of pearling result in fractions enriched in embryo and pericarp (F1), aleurone (F2) and subaleurone (F3). The iron concentration in the F1 fraction of TaVIT2 grain was decreased by 26% compared to control grain, in agreement with the percentage decrease in dissected embryos. It should be noted that the absolute iron concentration in the F1 pearling fraction is much lower than in dissected embryos, because of dilution by dry matter from the bran and the embryo at maturity. The F2 fraction contained 36.6% less iron in TaVIT2 grain, 49.8 ± 3.2 mg kg^−1^ dry weight compared to 77.7 ± 0.4 mg kg^−1^ in control grain (Fig. 1D). The inner starchy endosperm (core) contained ~2-fold more iron in TaVIT2 grain, whereas the iron concentration in the whole grain is similar in TaVIT2 and control grain, as noted previously (Connorton et al., 2017; Balk et al., 2019). The distribution of other metals such as Mn and Zn were little affected by TaVIT2 overexpression, except for small increases in the F1 fraction (Suppl. Fig. S3). The distribution of phosphorus in TaVIT2 grain was not affected, resulting in a lower P/Fe ratio in the core of the endosperm (Suppl. Fig. S3). Thus, overexpression of *TaVIT2* in the endosperm does not affect the amount of iron mobilized from the maternal plant into the grain, but does affect the distribution of iron within the grain: iron accumulates in a specific region of the endosperm at the expense of iron translocation to the embryo and aleurone.

### TaVIT2 endosperm cells accumulate iron in clusters of vesicles and globoids

Based on its function as a vacuolar iron transporter, overexpression of *TaVIT2* is expected to lead to iron accumulation in vacuoles. To obtain information on the subcellular location of iron in TaVIT2 grain, semi-thin transverse grain sections were stained with Perls’-DAB (Fig. 2). Dense granules of dark staining were observed in in the endosperm of TaVIT2 grain, but not in control grain (Fig. 2A, B). The region where intense iron staining was found corresponded with the blue Perls’ staining in hand-cut sections of TaVIT2 grain (Fig. 1A). Higher magnification images showed that the iron staining was confined to clusters of small round bodies of approximately 0.5 – 2 μm in diameter in the cytoplasm of endosperm cells (Fig. 2C; Suppl. Fig. S4). There appeared to be no association with other cell organelles, such as starch grains or the nucleus.

**Figure 2.**
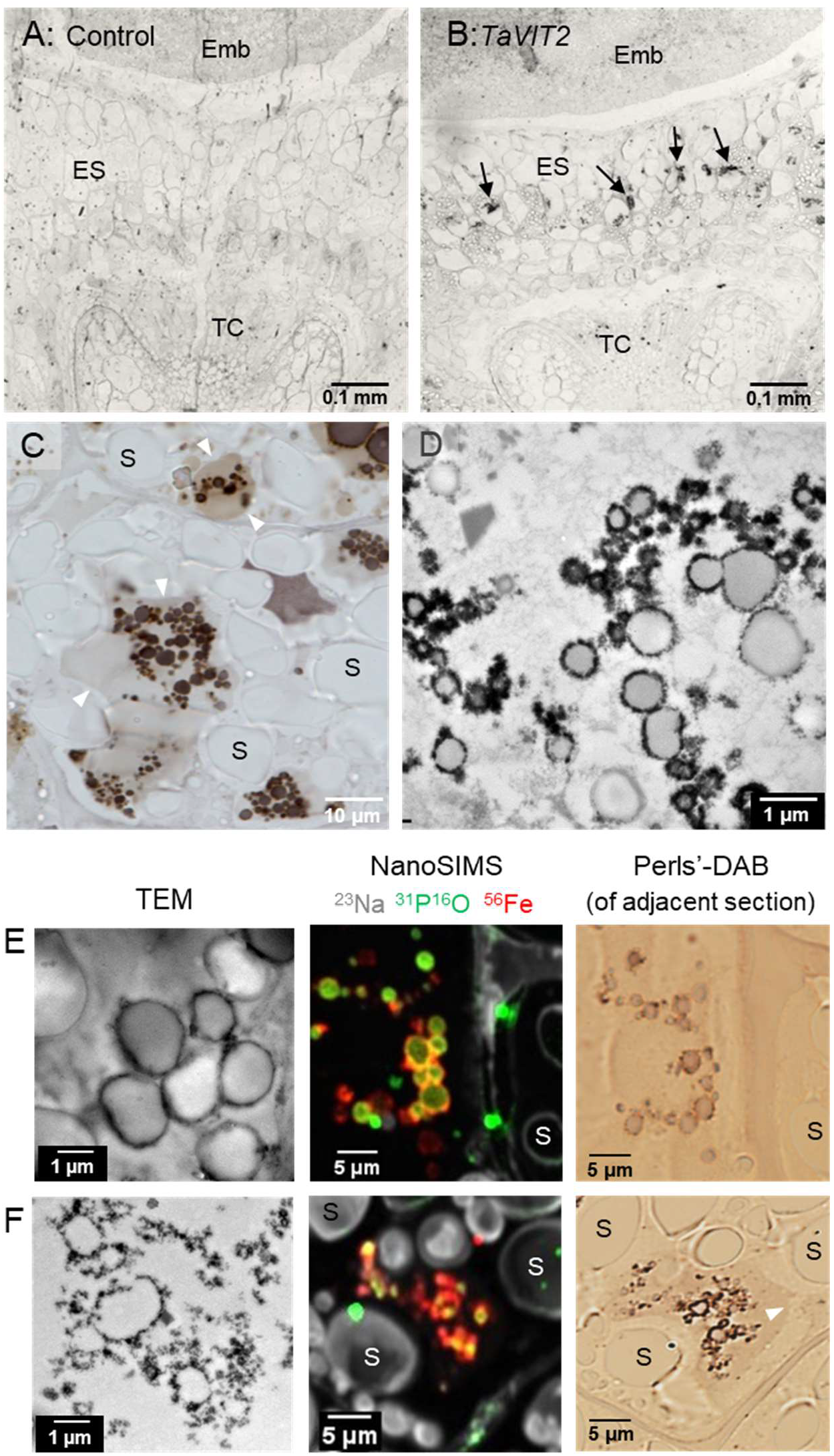
Iron accumulates in subcellular clusters in TaVIT2 endosperm cells. **A, B**. Transverse sections of wheat grains (21 dpa) from (A) control and (B) TaVIT2 stained for iron using the enhanced Perls’ method. Emb, embryo; Nuc, nucellar projection; TC, transfer cells of the nucellar projection. **C**. Detail of endosperm cells in TaVIT2 grain stained for iron, imaged with DIC microscopy. S, starch grain. **D**. Electron-dense structures in endosperm cells of TaVIT2 grain, imaged by TEM. **E, F**. Subcellular structures in (B) containing iron imaged by TEM (and EDS, see Fig. S5), NanoSIMS and Perls’-DAB staining. The NanoSIMS and Perls’-DAB images are from adjacent 1 μm sections.

Transmission electron microscopy (TEM) identified several distinct electron-dense morphological structures in the endosperm region of interest in TaVIT2 grain, which were absent from control grain. The most abundant morphologies were particles forming the outline of vesicles 0.2 – 0.8 μm in diameter but lacking any trace of a membrane (Fig. 2D, F); and clusters on the outside of membrane-bound vesicles 0.5 – 2.3 μm in diameter (Fig. 2E, Suppl. Fig. S5). Dispersed particles and aggregates of smaller particles were also observed but were less abundant (Suppl. Fig. S5A). Elemental analysis by Energy Dispersive X-ray Spectroscopy (EDS) indicated that all four electron-dense morphologies in TaVIT2 grain contained iron, whereas iron was not detectable outside these areas (Suppl. Fig. S5). The two most abundant types of iron-rich morphologies seen in TEM were also distinguishable by NanoSIMS (Fig. 2E, F). The NanoSIMS images were aligned with Perls’-DAB staining applied to adjacent sections. While the two morphologies looked identical with Perls’-DAB staining, this technique revealed that the smaller type are surrounded by an intracellular membrane (Fig. 2F, white arrow), indicating they are iron globoids inside a larger vacuole.

Accumulation of iron in a specific region of the starchy endosperm but not throughout, raises the question whether this coincides with a local abundance of iron or is due to localized expression of the *TaVIT2* transgene. The *TaVIT2* transgene is expressed using the *HMW Glu-1D×5* promoter, which is active in the entire starchy endosperm during grain filling as shown by promoter-GUS studies (Lamacchia et al., 2001). To verify that the expression pattern of *HMW-TaVIT2* is similar, we carried out in-situ hybridizations of 21 dpa grain. Hybridization with an antisense *TaVIT2* probe showed intense positive staining in all parts of the endosperm of TaVIT grain, especially in the subaleurone cells, matching the expected pattern of *HMW Glu-1D×5* promoter activity. In control grain, positive staining but of weaker intensity was seen in the aleurone cell layer, where endogenous *TaVIT2* is expressed. There was no signal in any tissue with the sense probe which served as negative control (Supplemental Fig. S6).

In summary, accumulation of iron-dense vesicles, representing iron trapped by overexpression of *TaVIT2*, indicates that the starchy endosperm region between the maternal transfer tissue and the embryo is a major transport route of iron in developing wheat grain.

### The rate of iron transport to the embryo is decreased in TaVIT2 grain

To study how iron is translocated into the developing grain at (sub)cellular resolution, we designed an experimental protocol for iron isotope labelling and subsequent NanoSIMS analysis (Fig. 3A). Iron-57 (^57^Fe) was chosen as a stable isotope over iron-54 and iron-58 because its mass differs sufficiently from other abundant elements in biological material and it has a relatively low natural abundance of 2.12%. By contrast, iron-54 has a natural abundance of 5.8% which would make it harder to detect enrichment after pulse-labelling. Although iron-58 has a very low natural abundance (0.28%) it is difficult to separate from nickel-58 (68.1% natural abundance) by mass. To detect iron by NanoSIMS, previously we used the caesium (Cs^+^) beam to detect FeO^−^(Moore et al., 2012), but the mass of ^57^Fe^16^O^−^ is nearly indistinguishable from the ^56^Fe^16^O^1^H^−^ion. However, the positive ions ^57^Fe^+^ (m/z = 56.9354) and ^56^Fe^1^H^+^ (m/z = 56.9428), generated with the oxygen (O^−^) beam, are sufficiently separated to allow reliable mapping of the ^57^Fe^+^ signal (Suppl. Fig. S7).

**Figure 3.**
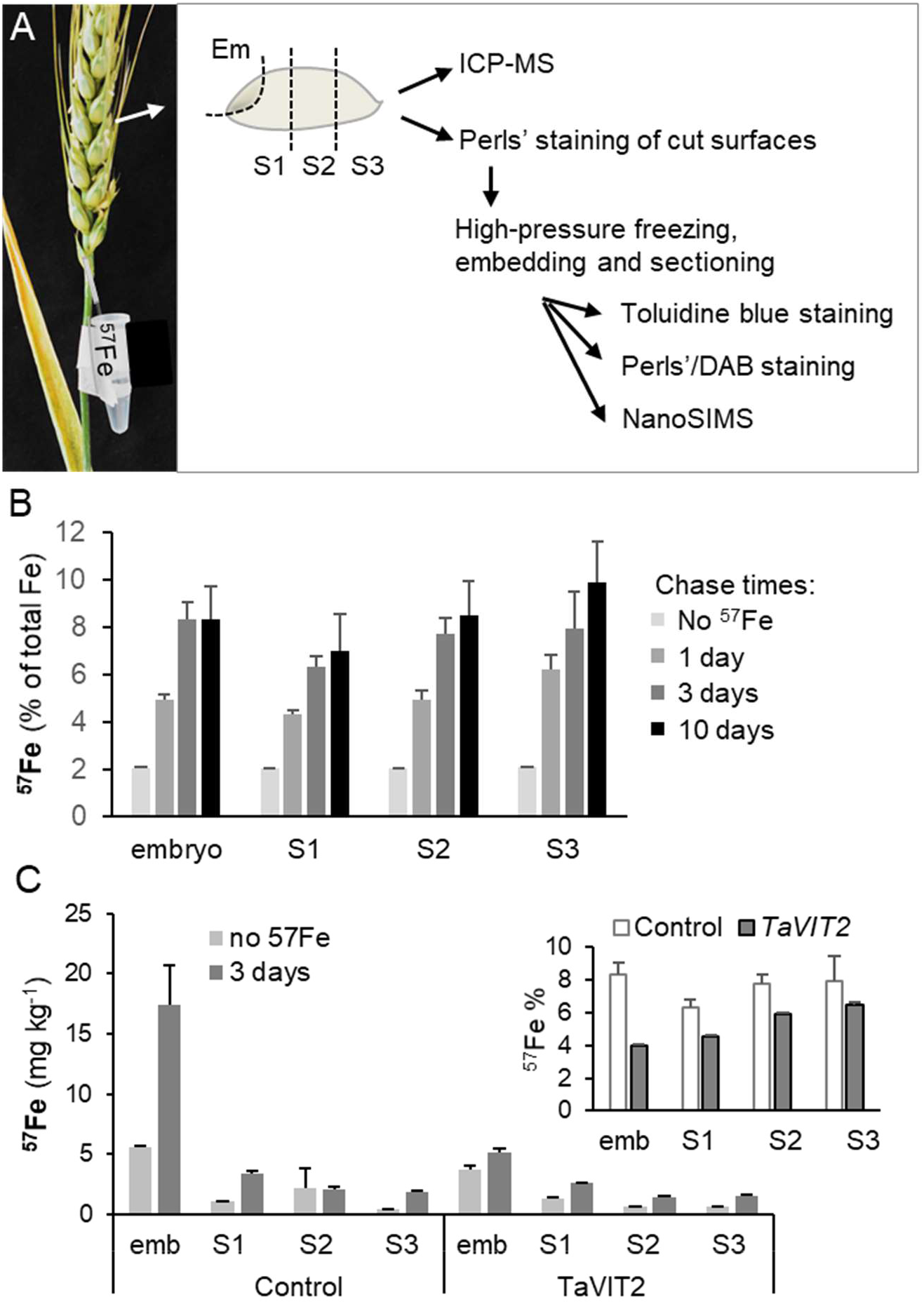
Iron isotope labelling of developing grain. **A**. Diagram of the experimental pipeline for ^57^Fe feeding and sample preparation. **B**. Percentage enrichment of ^57^Fe in parts of the grain as indicated in (A), following different chase times after pulse-labelling of 17-19 dpa wheat ears. Iron isotopes were quantified by ICP-MS. Values are the mean ± SE of 3 biological replicates, which are pools of 40-50 grain parts from one ear. **C**. Concentrations of ^57^Fe in the embryo (emb) and other parts of the grain (see A) in control and in plants overexpressing *TaVIT2* in the endosperm. ICP-MS values are given as absolute numbers and as percentage (inset), and are the mean ± SE of 3 replicates.

To directly target the iron isotope to the developing grains, a 1 ml solution of 50 μM ^57^Fe^3+^ and 0.5 mM citrate was fed into the base of the rachis (stem of the flowering spike) using a glass microcapillary tube (Fig. 3A). Feeding was performed on wheat ears around 18 days post anthesis (dpa), to coincide with the mid-stage of grain filling and iron mobilization (Waters et al., 2009; Beasley et al., 2019), and high activity of the *HMW* promoter driving expression of *TaVIT2* (Lamacchia et al., 2001). Following a 12 h isotope feeding pulse, the optimal chase time was determined experimentally, by ICP-MS measurement of ^57^Fe in the embryo and rest of the grain at 24 h, 72 h and 240 h (10 days), counting from the start of feeding. The relative abundance of ^57^Fe increased from background levels (2.12%) to ~5% after 24 h and to ~8% after 72 hours (Fig. 3B). Sampling the grain 10 days after isotope labelling showed no further increase in ^57^Fe enrichment, either because the grain filling period had come to an end or because non-labelled iron from the senescing leaves was mobilized at a sufficient rate to dilute the remaining isotope. Capillary insertion had no significant effect on grain development (Moore et al., 2016) and the total iron concentration in ‘fed’ and ‘non-fed’ samples was similar (data not shown).

The highest concentration of ^57^Fe accumulating over 72 h was in the embryos (Fig. 3C), indicating that most of the iron taken up at this stage is partitioned there. Interestingly, the increase in ^57^Fe enrichment was similar in proximal and distal parts of the grain, suggesting a constant rate of translocation to all grain tissues (Fig. 3C, inset). Overexpression of *TaVIT2* resulted in a marked decrease in ^57^Fe translocation to the embryo (by 2-fold), in agreement with the lower total iron concentration of TaVIT2 embryos (Fig. 1C). There was also a modest decrease in ^57^Fe translocation to the rest of the grain (Fig. 3C, inset). However, the ^57^Fe enrichment value in these parts is the sum of iron translocation into the endosperm and aleurone, two tissues with very different iron contents. To measure iron translocation into specific grain tissues, the ^57^Fe/^56^Fe was determined by NanoSIMS, averaging the counts for each isotope over 50 × 50 μm regions of interest (ROIs). Wheat samples harvested at 24 h after the start of isotope labelling showed a maximum of 5% enrichment of ^57^Fe compared to the natural ^57^Fe/^56^Fe abundance ratio of 2.31%, which was considered too little difference for quantification. After labelling for 72 h, up to 20% enrichment of ^57^Fe was found in ROIs.

^57^Fe and ^56^Fe counts from transfer cells, modified aleurone cells, the EAS region, central starchy endosperm, scutellum, embryo plumule and aleurone cells from 2 different biological replicates were collected and the average ratios are plotted in Fig. 4. In control grain, 12 – 18% ^57^Fe enrichment was found in transfer cells, modified aleurone, EAS and the embryo, but only 6 – 7% ^57^Fe enrichment in the starchy endosperm and outer aleurone cells. Most tissues in TaVIT2 grain showed a decrease in ^57^Fe/^56^Fe relative to control, with a small decrease in the maternal transfer cells and starchy endosperm, but a dramatic decrease in the EAS. At the time point of analysis (21 dpa), lots of iron has already accumulated in the EAS of TaVIT2 grain, and the low ^57^Fe/^56^Fe ratio indicates that the diversion into storage organelles has been saturated in this tissue.

**Figure 4.**
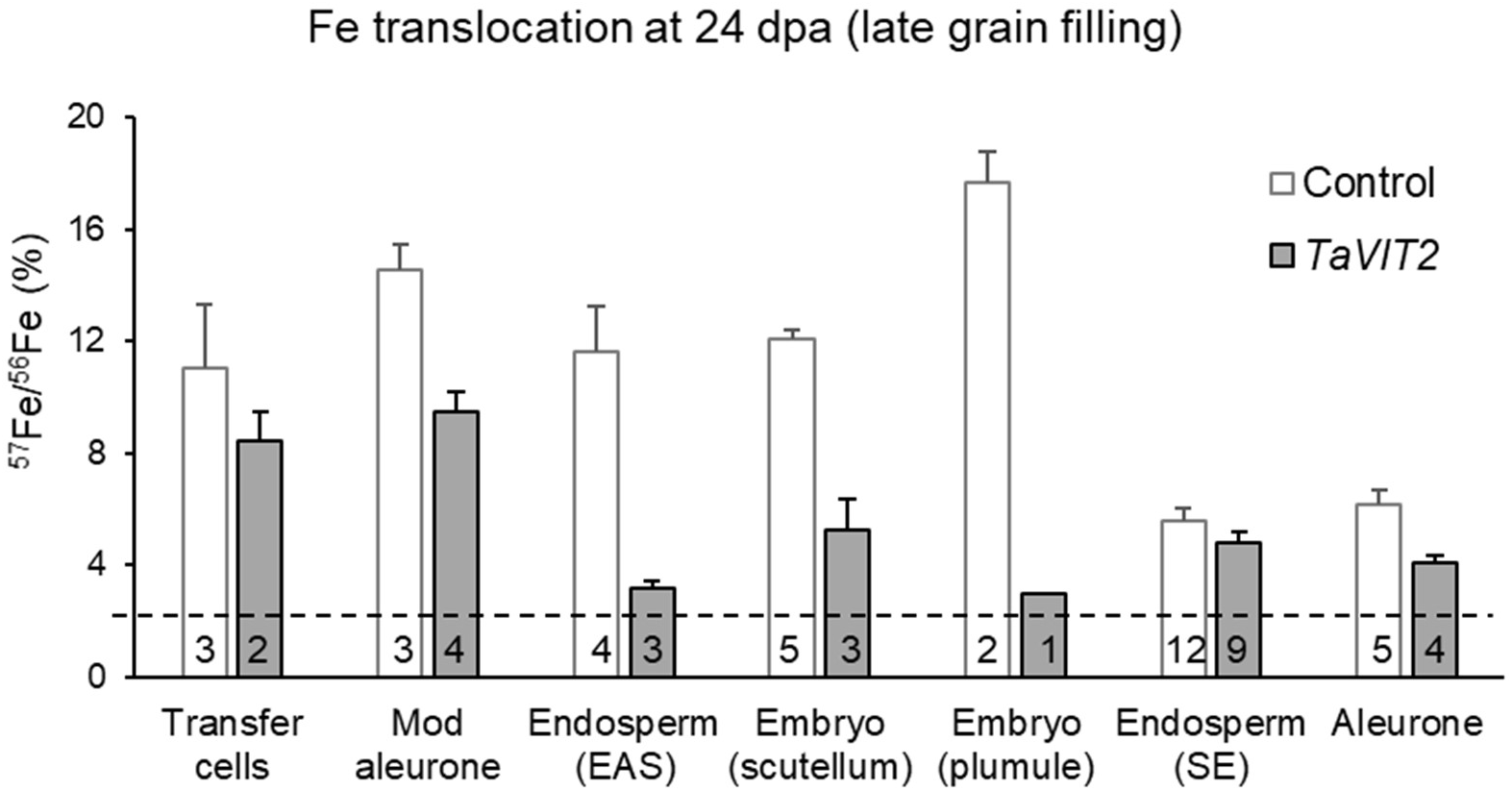
Iron translocation into different cell types following ^57^Fe labelling. Ratios of ^57^Fe and ^56^Fe signals determined by NanoSIMS, expressed as a percentage, from 50 × 50 μm regions of interest (ROIs) of different wheat grain tissues. The values are the mean ± SE of at least 2 ROIs in grain sections of 2 ^57^Fe-labelled wheat plants, except for the embryo plumule of TaVIT2 grain. The number of ROIs is given in each bar. The dashed line indicates the naturally occurring ^57^Fe/^56^Fe ratio (2.3%).

Taken together, ^57^Fe pulse labelling of developing wheat grain, analysed by ICP-MS in grain sectors or by NanoSIMS in specific tissues, therefore revealed a high flux of iron from the maternal transfer cells directly to the embryo, starting well before 21 dpa, which was perturbed by overexpression of *TaVIT2*.

### Iron is translocated in dynamic intracellular vesicles

Next, we exploited the combined methods of ^57^Fe pulse labelling and NanoSIMS to investigate iron dynamics within cells. Previous NanoSIMS analysis of ^56^Fe^16^O^−^ in durum wheat grains at 16 dpa showed that iron was concentrated in phosphate-containing globoids in the aleurone, whereas it was uniformly distributed in a starchy endosperm cell located 100 μm from the aleurone (Moore et al., 2012). Detection of ^56^Fe^+^ ions with the O^−^ beam showed a similar pattern of iron-containing globoids in the aleurone cells of bread wheat at 21 dpa (Fig. 5A; Suppl Fig. S8). Interestingly, ^57^Fe labelling revealed two populations of iron-dense structures, a population with 20 – 30% ^57^Fe enrichment 0.1 – 1 μm in diameter and a population with ~5% enrichment which were generally larger in diameter (0.5 – 2.5 μm) (Fig. 5B, Suppl. Fig. S8). The first population had virtually no associated PO signal, whereas the second population had a strong PO signal likely derived from phytic acid. In several instances, the ^57^Fe-rich structures are seen abutting larger globoids and were possibly merging (Fig. 5, white arrow heads). Overexpression of TaVIT2 in the endosperm led to a dramatic decrease in ^57^Fe entering the aleurone between 20 - 22 dpa, but the two population could still be recognised (Fig. 5C).

**Figure 5.**
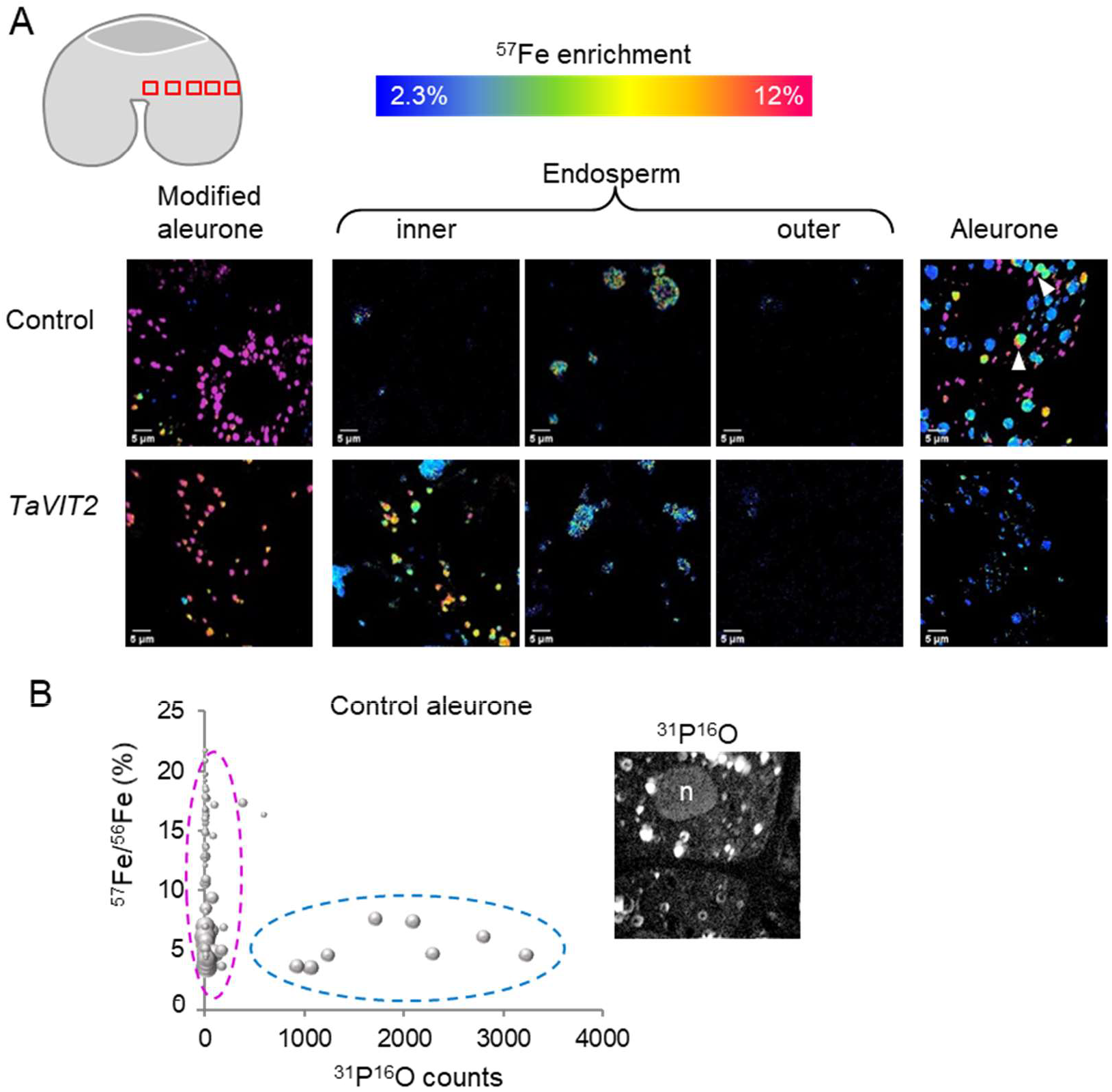
NanoSIMS analysis of ^57^Fe enrichment along the lateral axis in wheat grain. **A**. NanoSIMS scans (50 × 50 μm) of the indicated tissues in control and TaVIT grain. The position of the images is indicated in the cartoon image, top left. The ^57^Fe/^56^Fe ratio is represented by a colour scale from 2.3 – 12%. For the aleurone cells, the ^31^P^16^O signal is shown in the far-right image in grey scale. n, nucleus. **B** Relationship between ^57^Fe enrichment and ^31^P^16^O in iron-dense subcellular structures in aleurone cells of control grain. The size of each data point correlates with the area of the structure.

Transfer cells of the nucellar projection connect the vascular bundle from the mother plant to the grain, and thus play an important role in nutrient transport. The pattern of iron-rich bodies was comparable in transfer cells of control and TaVIT2 grain, showing a combination of disperse iron-rich vesicles and large clusters filling up most of the small cell volume (Fig. 6; Suppl. Fig. S9). The percentage ^57^Fe enrichment was decreased in transfer cells of TaVIT2 grain compared to control (Fig. 4 and 6), although this is a maternal tissue through which iron passes before it reaches the grain. However, compared to EAS region and embryo in TaVIT2 grain, ^57^Fe was significantly enriched.

**Figure 6.**
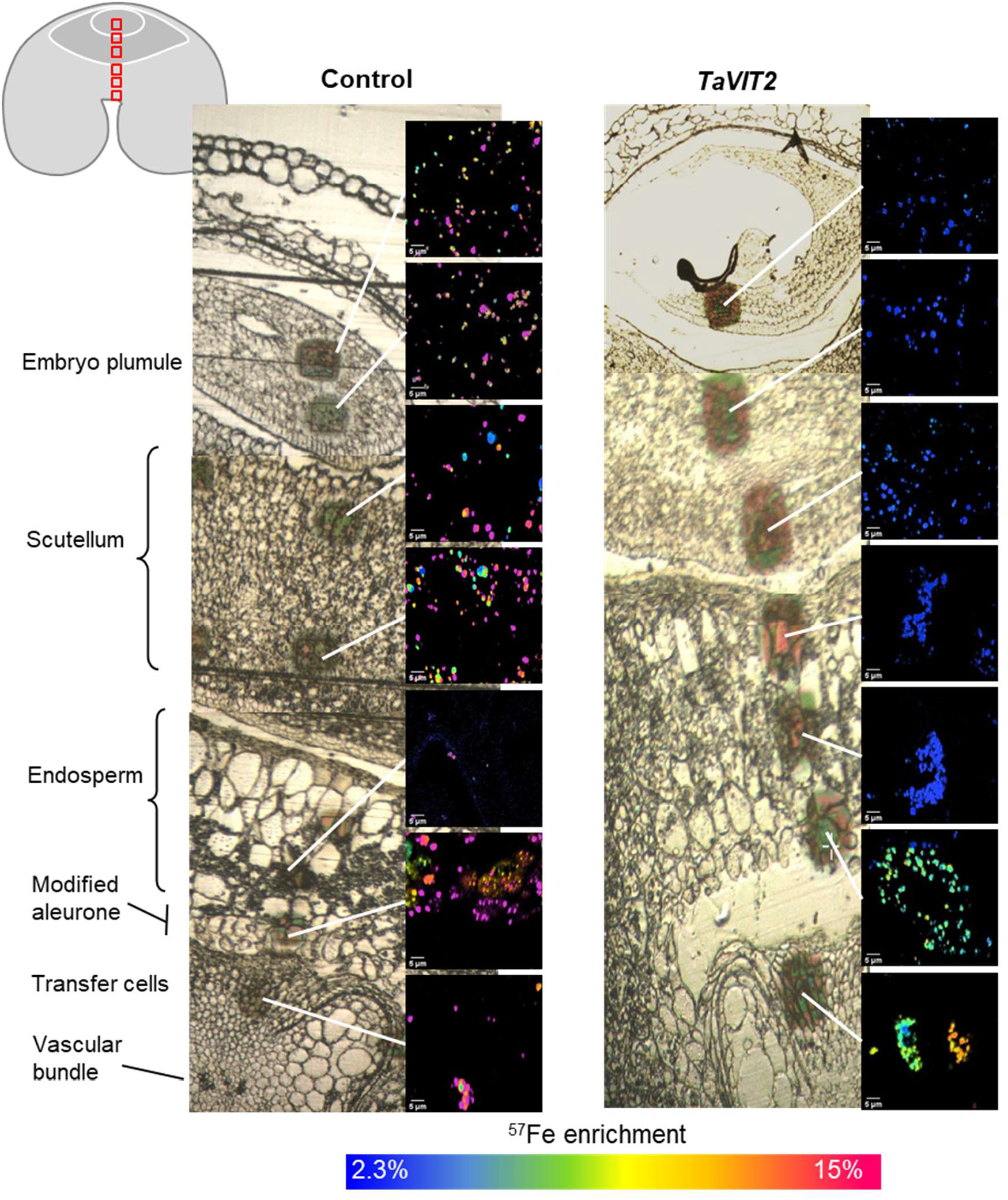
NanoSIMS analysis of ^57^Fe enrichment along the vertical axis in wheat grain. Light microscopy (LM) images of a transect through a control and a TaVIT grain with 50 × 50 μm images acquired by NanoSIMS showing the ^57^Fe/^56^Fe ratio. The position of the transect is indicated in the cartoon image, top left. The scan areas appear as dark patches in the LM image. The ^57^Fe enrichment is represented by a false colour scale from 2.3 – 15%.

In the modified aleurone (MA) cells adjacent to the transfer cells, the pattern of iron-rich bodies was different from that in aleurone cells around the periphery of the grain (Fig. 5A, 6). MA cells in both control and TaVIT2 grain showed an abundance of ^57^Fe-enriched vesicles, ranging from 6 – 20% enrichment in the control and 6 – 15% in TaVIT2. These patterns correlate with the specific function of MA cells, namely nutrient transfer rather than nutrient storage. Endosperm-specific expression of *TaVIT2* did have a negativeinfluence the iron dynamics in MA cells, intermediate to the suppression of iron translocation into the maternal transfer cells and EAS.

Starchy endosperm cells between the crease and embryo (EAS) contained only few iron-enriched vesicles 5 – 7 μm in diameter, whereas cells in the ‘cheeks’ of the developing grain showed a diffuse pattern of iron (Fig. 5A Fig. 6). In TaVIT2 grain, large clusters of ^56^Fe vesicles and globoids were found but with little or no enrichment in ^57^Fe. It is likely that the cells are saturated with iron prior to our time point of investigation and that iron transport into the cell is decreased. However, at the periphery of the EAS we observed a cluster of vesicles with smaller, ^57^Fe-rich vesicles on the outside and larger, Fe + PO containing vesicles towards the centre of the cluster (Fig. 7). Again, instances of abutting vesicles suggest that fusion is taking place, with ‘newer’ ^57^Fe being delivered to ‘older’ ^56^Fe stored with PO.

**Figure 7.**
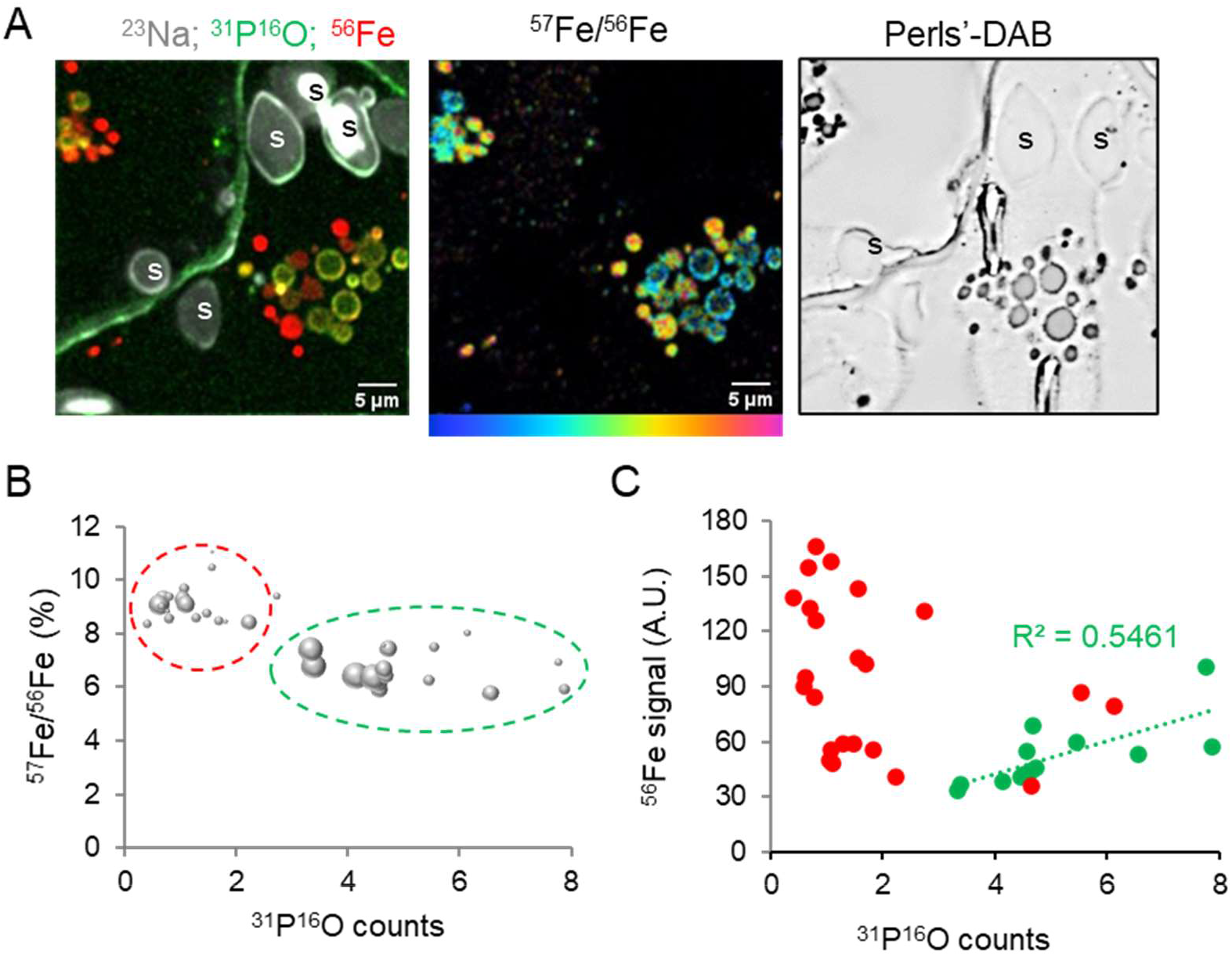
Heterogenous composition of iron-rich vesicles. **A**. NanoSIMS images of endosperm cells at the periphery of the EAS in TaVIT2 grain, showing the distribution of ^23^Na, ^31^P^16^O and ^56^Fe (left) and the ^57^Fe/^56^Fe ratio (middle). The same area in an adjacent 1 μm section stained with Perls’-DAB is shown on the right. A cell wall runs through the middle of the image. s, starch grain. **B**. Relationship between the ^57^Fe enrichment and ^31^P^16^O signal in iron-rich vesicles. The size of each data point corresponds to the area of the vesicle. **C**. Relationship between the ^56^Fe and ^31^P^16^O signal from the iron-rich vesicles.

The scutellum contained a large number of iron-rich, membrane-bound vesicles (Fig. 6), in agreement with iron accumulation seen by XRF (Neal et al., 2013; Singh et al., 2013; De Brier et al., 2016). Cells in the outer cell layers of the scutellum, adjacent to endosperm cells that are undergoing programmed cell death, tended to have a higher enrichment with ^57^Fe. The plumule of the embryo also showed a high density of iron-rich vesicles. In the TaVIT2 overexpressing grain, the density of the iron vesicles was similar to control grain (Fig. 6), but ^57^Fe enrichment was dramatically decreased (Fig. 4), in agreement with the lower total iron content of TaVIT2 embryos (Fig. 3C).

In summary, the presence of iron-rich organelles in most cell types (except the starchy endosperm) and their heterogeneity in ^57^Fe enrichment suggest that vesicle-mediated transport is a major route of cell-to-cell iron translocation.

## DISCUSSION

To gain insight into iron translocation and distribution into developing cereal grains, we developed a protocol for ^57^Fe labelling combined with NanoSIMS. This revealed how different cell types contribute to iron translocation with remarkable differences in their cell biology. A summary of the findings is presented in Fig. 8. The main flux of iron is from the maternal transfer cells to the embryo through a specific zone of the starchy endosperm, as shown by ^57^Fe enrichment in embryos of control grain (Fig. 3C) and entrapment of this iron flux when TaVIT2 is overexpressed under the control of an endosperm-specific promoter (Figs. 1-4). A role in nutrient transport for this part of the endosperm, previously termed Endosperm Adjacent to Scutellum (EAS), has been suggested based on transcriptomics analysis in maize, which showed a marked upregulation in this tissue of transporters for sugars, amino acids and some metal transporters (Doll et al., 2020). The amount of iron travelling to the aleurone layer at the periphery of the endosperm was also strongly affected by *TaVIT2* overexpression (Fig. 4 and 5), although no iron accumulation was observed in tissues adjacent to the aleurone, and the route of this pool of iron therefore remains unclear.

**Figure 8.**
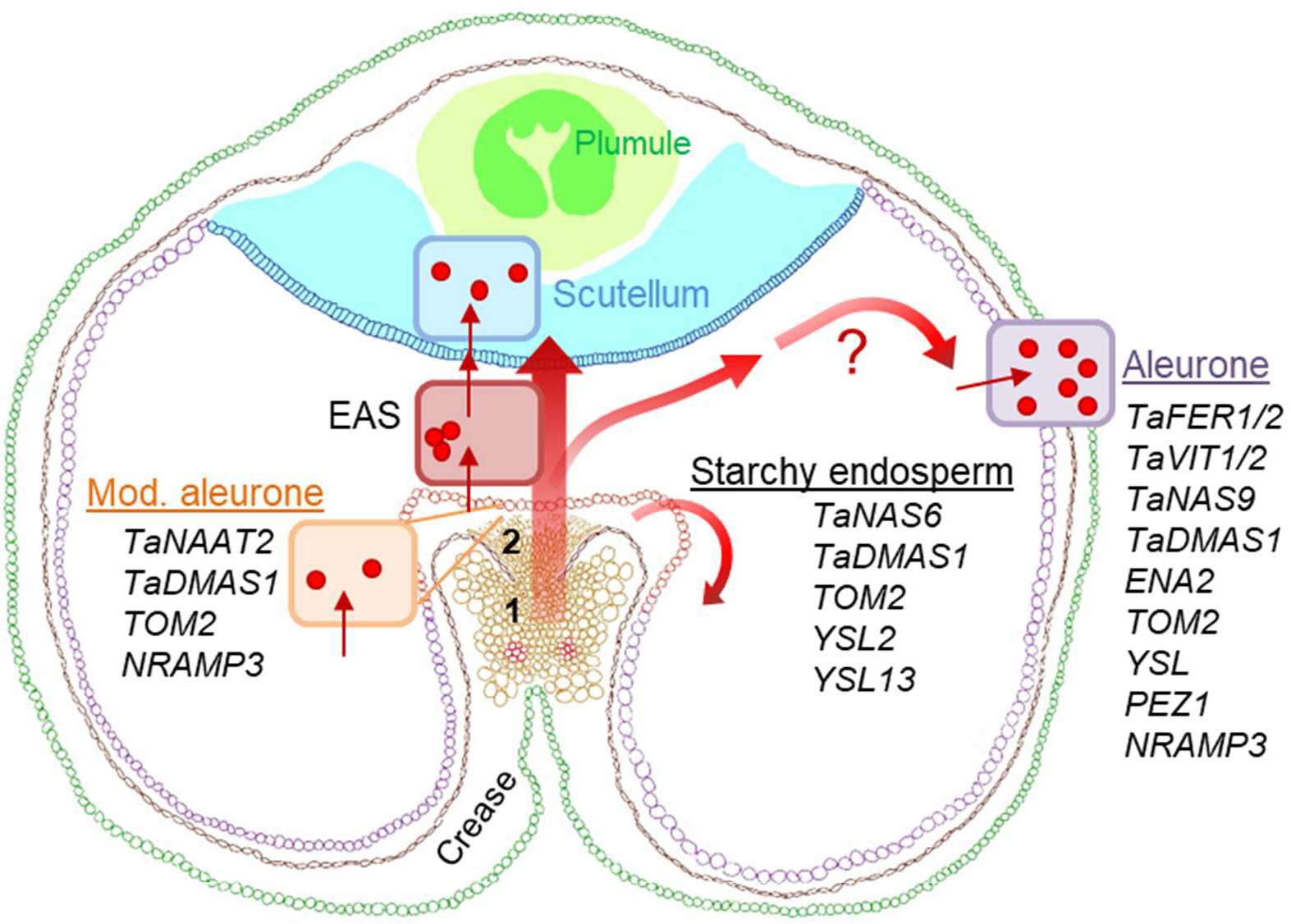
Proposed model for iron translocation in developing wheat grain. Iron enters the developing grain from the maternal vascular bundle (1) running along the ventral crease, through the transfer cells of the nucellar projection (2) and the modified aleurone. The main flux of iron is directly to the scutellum of the embryo, via the Endosperm Adjacent to the Scutellum (EAS). Transcripts of iron homeostasis genes that are relatively abundant in either the modified aleurone, starchy endosperm and/or aleurone based on RNA-seq data from Pfeifer et al., (2014) are indicated. *TaXXX* indicates a gene annotated in wheat, *Triticum aestivum*, otherwise the rice acronyms are used.

NanoSIMS analysis of ^57^Fe provided subcellular detail of how iron is trafficked through the cell by vesicles and vacuolar globoids. Subcellular imaging of ^57^Fe was facilitated by the recent addition of the RF O- source to the NanoSIMS allowing measurement of positively charged iron ions with discrimination of ^57^Fe^+^ from ^56^FeH^+^. By contrast, the negative iron ions could not be resolved with the Cs+ beam (Fig. S7). The combination of isotope labelling and NanoSIMS is a powerful technique to study dynamic transport processes, including ^15^N/protein accumulation in wheat grain (Moore et al., 2016), dimethylsulfoniopropionate metabolism using ^34^S in microalgae (Raina et al., 2017) and ^70^Zn uptake in radish roots (Ondrasek et al., 2019). Although NanoSIMS cannot determine the precise concentration of ^57^Fe, the enrichment of ^57^Fe compared to ^56^Fe with similar physical properties can be reliably determined, as shown by the small statistical variance between different images of the same tissue and biological replicates (Fig. 4). While our analysis was restricted to one time point in grain development (21 dpa), it nevertheless provided a dynamic snapshot of iron translocation by visualizing the ^57^Fe/^56^Fe ratio and distinguishing between ‘old’ iron and newly translocated iron.

The small, highly ^57^Fe-enriched bodies in the aleurone cells (Fig. 5A) have not previously been observed. The ^57^Fe enrichment indicates that this is a pool of iron that has recently entered the cell. A previous analysis of the aleurone with the Cs+ beam showed the larger iron-phytate globoids (Moore et al., 2012). These have a much lower ^57^Fe/^56^Fe ratio and thus contain mostly iron present before the start of isotope labelling. It is probably that the higher sensitivity achieved by mapping iron as Fe^+^ with the O^−^ beam, rather than FeO^−^ with the Cs^+^ beam, enabled visualization of these iron particles, which are low in PO, and thus contain little phosphate or phytic acid (Fig 5B). Because free iron is cytotoxic, iron in these small bodies may be bound to citrate or nicotianamine (NA). A pool of Fe-citrate was found in aleurone cells of mature wheat grains using XANES imaging (De Brier et al., 2016; Pongrac et al., 2020). These studies of mature grains failed to identify Fe-NA in aleurone cells and this iron species may only occur during grain filling. At 20 dpa, transcripts of *TaNAS9*, encoding a NA synthase, are abundant in aleurone cells (Suppl. Fig. S10), which is approximately the same stage as our NanoSIMS analysis. Other techniques such as Time-of-Flight SIMS analysis of small organic molecules could be used to establish if NA is present in younger grain tissues.

Detection of vesicles enriched in ^57^Fe in all grain tissues from both control and TaVIT2 grain indicates that iron translocation is predominantly symplastic. Only a small number of NanoSIMS images captured an iron signal in cell walls, see Suppl. Fig S11. It is currently unclear how iron is transported through the plant cytosol, although NA is thought to play a key role (Mari et al., 2020). Our study suggests that vesicle transport could also be involved. Vesicle-mediated transport of iron has been reported in erythrocytes, where a large flux of iron is delivered to the mitochondria for haem biosynthesis (Hamdi et al., 2016).

The observed cell-type specific patterns of iron uptake and storage are likely to be underpinned by differential gene expression. An RNA-seq study by Pfeifer and colleagues (2014) investigated gene expression in tissue samples enriched in modified aleurone cells, starchy endosperm and aleurone cells of wheat grains at 20 dpa. We reanalysed this data set to extract the expression levels of wheat genes likely to be involved in iron homeostasis, based on sequence homology to genes in rice and Arabidopsis (Suppl. Fig. S10). Ferritin and *VIT* genes were noticeably upregulated in the aleurone tissue, but not in the modified aleurone or starchy endosperm, in agreement with our NanoSIMS data (Fig. 5). NA is known to be important for iron loading into the grain (Mari et al., 2020). Interestingly, two different NAS genes were upregulated in aleurone and starchy endosperm, *TaNAS9* and *TaNAS6*, respectively. *TaNAS9* is orthologous to *OsNAS3* (Bonneau et al., 2016), but there is no direct orthologue of *TaNAS6* in rice. Enzymes encoded by *NAAT* and *DMAS* work sequentially to convert NA to a 3”-oxo intermediate and then to deoxymugineic acid (Beasley et al., 2017). Transcripts of the *TaNAAT2* triad are enriched in the modified aleurone but not in the other two tissues. This suggests that the modified aleurone converts NA to the 3”-oxo intermediate, which could be further converted to deoxymugineic acid by *TaDMAS1* expressed in all three tissues. Wheat homologues of the mugineic acid exporter *OsTOM2* are also expressed in all three tissues, but the homologue of the NA exporter *OsENA2* is primarily expressed in aleurone cells.

Elevated aleurone expression of a wheat gene triad related to *OsPEZ1*, encoding a phenolics exporter active in the xylem (Ishimaru et al., 2011), suggests that enhancing the solubility of iron in cell walls may facilitate iron uptake as well. Uptake of Fe-NA complexes is mediated by YSL transporters, of which seven different paralogues are expressed in the investigated wheat grain tissues. The Fe-NA transporter ZmYSL2 was recently shown to be important for iron import into the embryo and aleurone cells in maize kernels (Zang et al., 2020), and this gene may correspond to *TraesCS2D02G387800* in wheat (Suppl. Fig. S10C). Transcripts of the wheat homologues of *OsYSL6* and *OsYSL9* are also enriched in aleurone cells. Interestingly, a rice mutant line of *OsYSL9* had decreased amounts of iron in the embryo, but increased contents in the polished grain (mostly endosperm), indicating that OsYSL9 plays a role in iron translocation from the EAS to the embryo (Senoura et al., 2017).

Dynamic iron studies, gene expression data and functional genetics will be invaluable for developing new biofortification strategies. Our isotope labelling study indicated that endosperm-specific expression of *TaVIT2* was successful as a biofortification strategy because iron is captured from the large flux going through the EAS and retained there, starting before 20 dpa (Fig. 3 and 4). Interestingly, previous studies overexpressing ferritin under the same promoter did not lead to accumulation of significant amounts of iron in the endosperm (Neal et al., 2013). A possible reason for this difference is that ferritin is a facultative iron store, which can release iron as easily as taking it up. In TaVIT2 grain, iron accumulated in a relatively small part of the endosperm, and the challenge will be to direct more iron into the ‘cheeks’ of the grain. Moreover, increasing total grain iron, rather than redistributing it, would be necessary. It was recently shown that the latter can be achieved by overexpressing the rice NAS2 gene under the constitutive *UBIQUITIN* promoter (Beasley et al., 2019). Because of different modes of action of the *OsNAS2* and *TaVIT2*, combining these two transgenes is likely to have a synergistic effect, increasing iron in wholemeal and white flour fractions simultaneously. Moreover, higher NA levels should also raise the concentration of zinc and lead to higher bioavailability of both micronutrients (Beasley et al., 2019). Other genes, such as those encoding grain-specific YSL and PEZ transporters, may also be interesting candidates for biofortification. Isotope labelling and imaging can make an important contribution in understanding cell-specific processes of iron homeostasis, helping to inform such biofortification strategies.

## MATERIALS AND METHODS

### Plant material and growth

Transformation of wheat lines (*Triticum aestivum* var. Fielder) with pBract202-TaVIT2 and their initial characterization have been described in (Connorton et al., 2017; Hayta et al., 2019). As a control, we used the offspring of a plant regenerated after transformation that tested negative for the transgene. This line, 22-15, did not accumulate iron in the endosperm and white flour fraction (Table S1; Connorton et al., 2017). All analyses were carried out using the T3 generation, except for the germination tests for which T4 grain was used. Plants were grown in a glasshouse kept at approximately 20°C with 16 h of light. Plants were watered as required.

### T-DNA copy number and TAIL-PCR

To estimate the numbers of T-DNA copies in individual plants, quantitative real time PCR analysis was carried out similar to the approach taken by (Bartlett et al., 2008) using DNA from seedlings of the T4 generation. Thermal asymmetric interlaced polymerase chain reaction was performed essentially as described in (Wu et al., 2015), except that recombinant Taq polymerase purified from Escherichia coli was used (Engelke et al., 1990).

### Element analysis by ICP-OES and ICP-MS

Preparation of flour fractions and element analysis were essentially carried out as previously described (Tosi et al., 2011; Connorton et al., 2017). See Supplemental Methods for details.

### Isotope labelling and sample preparation for microscopy

Iron-57 in the form of ^57^Fe_2_O_3_ (Cambridge Isotope Laboratories, Tewksbury, MA, USA) was added to a small volume of concentrated hydrochloric acid and incubated at 37°C overnight until it was dissolved. The solution was diluted with H_2_O to obtain 20 mM ^57^Fe in 1 M HCl and stored at 4°C. The ^57^Fe feeding solution (50 μM ^57^Fe, 0.5 mM Na citrate and 10 mM MES-KOH pH 6.0) was freshly prepared by mixing 25 μl ^57^Fe-HCl stock solution, 50 μl 100 mM Na citrate, 10 μl 1 M NaOH and 1 ml 100 mM MES buffer, made up to a final volume of 10 ml with H_2_O. After adjusting the pH to 6.0, 1 ml was filtered (pore size 0.22 μm) and pipetted into a 1.5-ml Eppendorf tube attached to the ear with tape. A 10 μl glass microcapillary tube (Drummond, Sigma Aldrich) was placed in the tube and the other end was inserted into the rachis. The solution was taken up naturally by the wheat ear over the course of ~12 h. The grains (40-50) from one ear were harvested after 24, 72 or 240 h and dissected into four parts: embryo, and three equal remaining sectors (S1, S2 and S3), and then were freeze-dried for ICP-MS. Four grains from the middle of the ear were harvested for high pressure freezing, sectioning and microscopy.

Transverse slices of the grain (0.1 mm thick, see Fig. 3 for positions) were infiltrated with 0.5 M MES-KOH pH 5.4, and frozen using a high-pressure freezer (HPM 100 from Leica Microsystems, UK). Freeze substitution, embedding in LR White resin (Agar Scientific UK, R1281) and sectioning was as described in Moore et al., 2012; 2016).

### Iron staining using the Perls’ method

Iron staining of tissue was performed as described in (Meguro et al., 2007; Roschzttardtz et al., 2009) using the Perls’ method, enhanced with 3,3’-diaminebenzidine where indicated.

### TEM-EDS analysis

The embedded wheat grain samples used for NanoSIMS were sectioned with a Leica UC7 ultramicrotome (Leica Microsystems, Vienna, Austria), and 100-nm ultrathin sections were mounted on 200-mesh copper TEM grids (Agar scientific). Where indicated, sections were contrasted with 2% uranyl acetate for 10 min, and Reynold’s lead citrate (Reynolds, 1963) for 2 min. TEM imaging was done with a 200kV JEOL 2011 TEM (JEOL Ltd., Tokyo, Japan). Circular regions of interest were chosen for energy dispersive x-ray spectroscopy (EDS) analysis. X-rays were detected with an Oxford INCAx-sight detector (Oxford Instruments, Abingdon, UK), and analyzed with the Oxford INCA Suite version 4.02.

### NanoSIMS

NanoSIMS analysis was performed with a Cameca NanoSIMS 50L (CAMECA, France), explained in detail elsewhere (Hoppe et al., 2013). A 16 keV O^−^ ion beam with a current between 45-85 pA and a beam size of approximately 600 nm (L1 = 1300-1582, D1 = 2 (300 μm aperture) was focused onto the sample and rastered over the surface to generate positive secondary ions. The L1 lens was used to increase the beam current and counts of ^57^Fe, and reduce analysis time, but this resulted in a lower spatial resolution than can typically be achieved with the NanoSIMS (100 nm). The entrance slit was set to position 3 (20 μm width), and the aperture slit to position 2 (200 μm width).

The detectors were aligned to simultaneously detect ^23^Na^+^, ^31^P^+^, ^40^Ca^+^, ^31^P^16^O^+^, ^56^Fe^+^ and ^57^Fe^+^. Detectors were aligned using bulk standards and particular care was taken to minimise the mass interference of ^56^Fe^1^H^+^ with ^57^Fe^+^ (Suppl. Fig. S7). Prior to analysis of regions of interest, a depth profile was acquired from a bulk Fe standard to check the measured ^57^Fe/^56^Fe isotope ratio and compare it to the natural isotope ratio of 2.3%. The detector for ^57^Fe was set slightly to the left of the centre of the peak to avoid the mass interference of ^56^Fe^1^H^+^. This resulted in a slightly lower measured isotope ratio than natural, with an average of 2.24% across all measurements (Suppl. Fig. S7C). All regions of interest and subcellular features were however enriched by significantly more than the 0.06% difference in the ratio.

Regions of interest were selected using the charge-coupled device (CCD) camera on the NanoSIMS. The samples were coated with 40 nm of platinum (Pt) prior to loading into the instrument to minimise sample charging. The Pt was removed by repeatedly scanning a defocused O^−^ beam (D1=0) over an area of 70 × 70 μm with a total dose of 2.55 – 3 × 10^17^ ions/cm^−2^. Following implantation, ion images were acquired using a focused beam over an area of 50 × 50 μm with 256 × 256 pixels and dwell time of 2000 μm per pixel. Several hundred images of each region of interest were acquired and summed together to improve the statistics.

Data processing was conducted with FIJI software using the OpenMIMS plugin (Harvard, Cambridge, MA, USA). Image processing included drift correction, summing of images, selecting regions of interest (ROIs) and extracting counts from them and generating colour merge and Hue Saturation Intensity (HSI) ratio images to show isotopic variation of ^57^Fe/^56^Fe.

### Bioinformatics

Iron homeostasis genes in wheat were taken from (Borrill et al., 2014) and supplemented with literature searches to a total of 232 genes including homeologues. Publicly available RNA-seq datasets of MA, SE and AL-enriched samples (Pfeifer et al., 2014) were mapped to the IWGSC RefSeq v1.1 gene models using Kallisto v0.43.1 (Bray et al., 2016). Gene expression was analyzed using the R package Sleuth v0.30.0 with default settings. This identified 213 transcripts from 67 wheat iron homeostasis genes that were expressed in at least one tissue. Transcript Per Million (TPM) means ± SD were calculated from 4 replicates in the RNA-seq data set. To filter for real expression over noise, a cut-off of 1 TPM in at least one tissue was imposed, and a q-value of significance <0.05 as calculated using the Likelihood Ratio Test.

## Acknowledgements

We would like to thank Kirstie Halsey (Rothamsted Research) for sample preparation for microscopy; Mark Durenkamp (Rothamsted Research) and Graham Chilvers (University of East Anglia) for ICP-OES and ICP-MS analyses; Eva Wegel (John Innes Centre) for microscopy assistance; Sophie Harrington and Jemima Brinton (John Innes Centre) for help with bioinformatics and Kexue Li for help with NanoSIMS. This research was funded by the Biotechnology and Biological Sciences Research Council, grant awards BB/P019072/1 (S.S., Y.W., J.M.C., P.R.S., K.L.M. and J.B.) and BB/T004363/1 (Q.X.), and a Newton Fellowship NF171396 to S.K.V. The NanoSIMS was funded by UK Research Partnership Investment Funding (UKRPIF) Manchester RPIF Round 2. This work was also supported by the Henry Royce Institute for Advanced Materials, funded through EPSRC grants EP/R00661X/1, EP/S019367/1, EP/P025021/1 and EP/P025498/1.

## Data Availability

The data that support the findings of this study are available in the supplementary material of this article and, if not there, available from the corresponding author upon reasonable request.

## Author Contributions

SS, YW, EV, SKV, QX, JW and JRC performed experiments and analysed data. JRC, PRS, KLM and JB planned the research. KLM and JB wrote the manuscript.

---

**Supplemental Table S1.** Copy number analysis and iron concentration in white flour of *HMW-TaVIT2* wheat lines.

|  | Line | Fe (mg/kg)<br>in white flour | Zygosity<sup>a</sup> | T-DNA<br>copies<sup>b</sup> |
| --- | --- | --- | --- | --- |
| T<sub>0</sub> | 22-19 | 18.7 ± 0.9 | het | 8 |
| T<sub>1</sub> | 22-19-1 | ND | het | 6 |
|  | 22-19-2 | ND | wt | 4 |
|  | 22-19-3 | 17.3 ± 1.4 | het | 4 |
|  | 22-19-4 | 19.9 ± 0 | **homo** | 8 |
|  | 22-19-5 | 13 ± 0.1 | het | 8 |
|  | 22-19-6 | 15.6 ± 0.5 | wt | 2 |
|  | 22-19-7 | 17.6 ± 0.3 | het | 10 |
|  | 22-19-8 | 12.2 ± 0.9 | wt | 2 (4) |
|  | 22-19-9 | 10.8 ± 0.1 | wt | 6 |
|  | 22-19-10 | 14.6 ± 0 | **homo** | 8 |
|  | 22-19-11 | 18.7 ± 0.8 | het | 6 |
|  | 22-19-12 | 15.1 ± 0 | wt | 6 |
| T<sub>1</sub> | 22-29-7 | 9.6 ± 0.3 |  | 0 |
|  | 22-29-12 | 8.9 ± 0.2 |  | 0 |
| T<sub>1</sub> | 22-15-12 | 7.4 ± 0 |  | 0 |

| T<sub>2</sub> line | T-DNA<br>copies<sup>b</sup> |
| --- | --- |
| 22-19-4-1 | 9 |
| 22-19-4-2 | 8 |
| 22-19-4-3 | 10 |
| 22-19-4-4 | 10 |
| 22-19-4-5 | 9 |
| 22-19-4-6 | 8 |
| 22-19-4-7 | 11 |
| 22-19-4-8 | 10 |
| 22-19-4-9 | 11 |
| 22-19-4-10 | 8 |
| 22-19-4-11 | 10 |

<sup>a</sup>Zygosity for the T-DNA insertion in *TraesCS4D02G046700* (*TaUEV1A-4D*) determined by PCR. het, heterozygous; wt, wild type lacking a T-DNA insertion; homo, homozygous.
<sup>b</sup>The number of T-DNA copies was determined by qPCR using primers for the hygromycin selectable marker gene. Higher values (>4) are approximate because of the log scale of the data output.

**Supplemental Figure S1.**
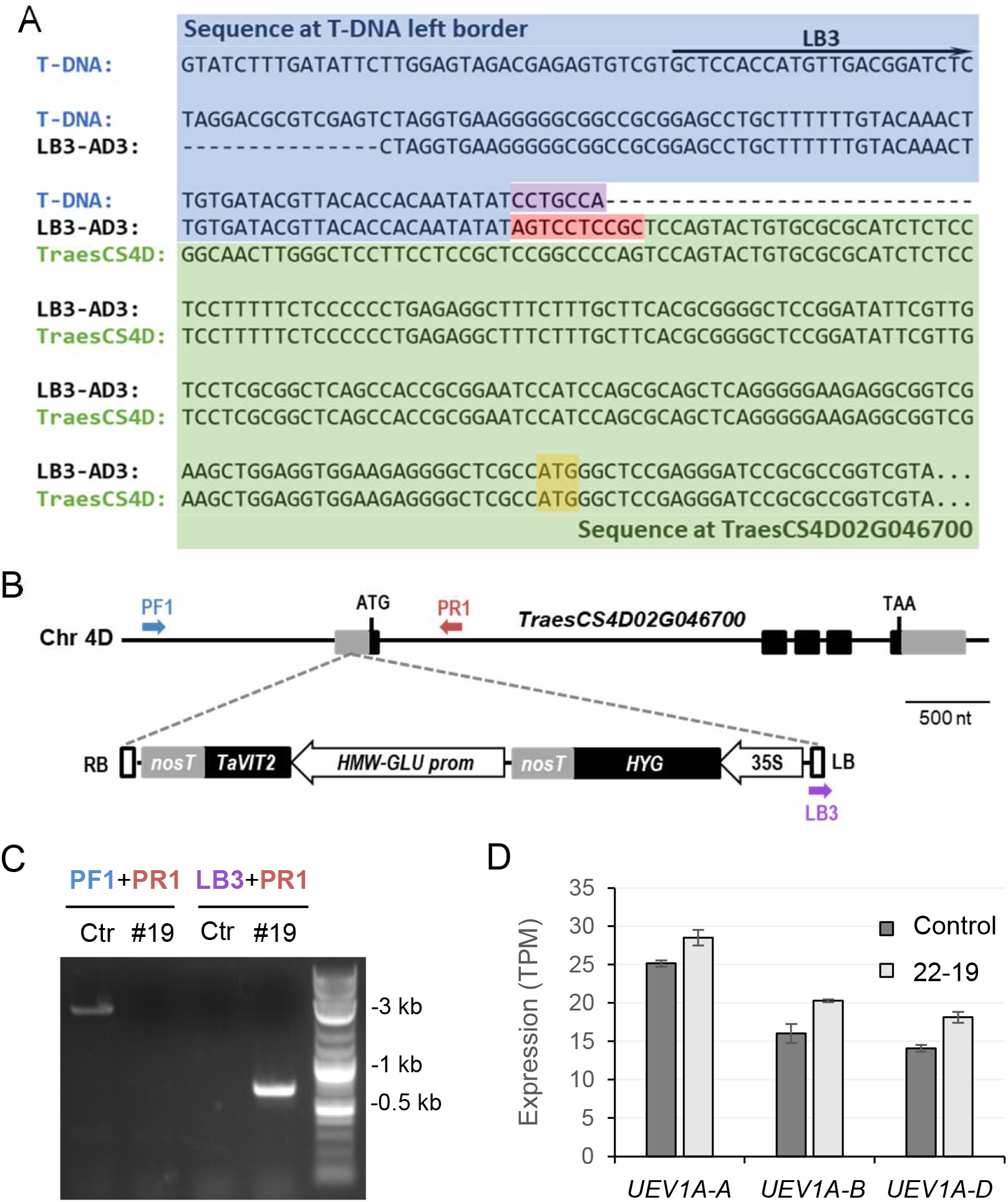
Insertion site of the T-DNA in line 22-19. **A**. Sequence of the TAIL-PCR product resulting from the left border (LB)-specific primers and arbitrary degenerate primer AD3, aligned with the T-DNA LB sequence (blue) and wheat genomic sequence matching the 5’-UTR of TraesCS4D02G046700 (green). There is a 7-bp truncation of the T-DNA left border (purple) and a 10-bp insertion (red). The start codon is marked yellow. **B**. Diagram of the T-DNA insertion into chromosome 4D. RB, right border; LB, left border; nosT, nopaline synthase terminator; TaVIT2, wheat VIT2 gene; HMW-GLU prom, High Molecular Weight glutenin promoter; HYG, hygromycin resistance; 35S, CaMV 35S promoter; PF1, PR1 and LB3 are primers. **C**. Verification of the T-DNA insertion site by PCR using the primers indicated in (B) **D**. Expression of *TraesCS4D02G046700* (TaUEV1A) homeologs in developing grain 21 days post anthesis of control and line 22-19 from RNA-seq data. Values are the average of 3 biological samples, with error bars representing SE of the mean.

**Supplemental Figure S2.**
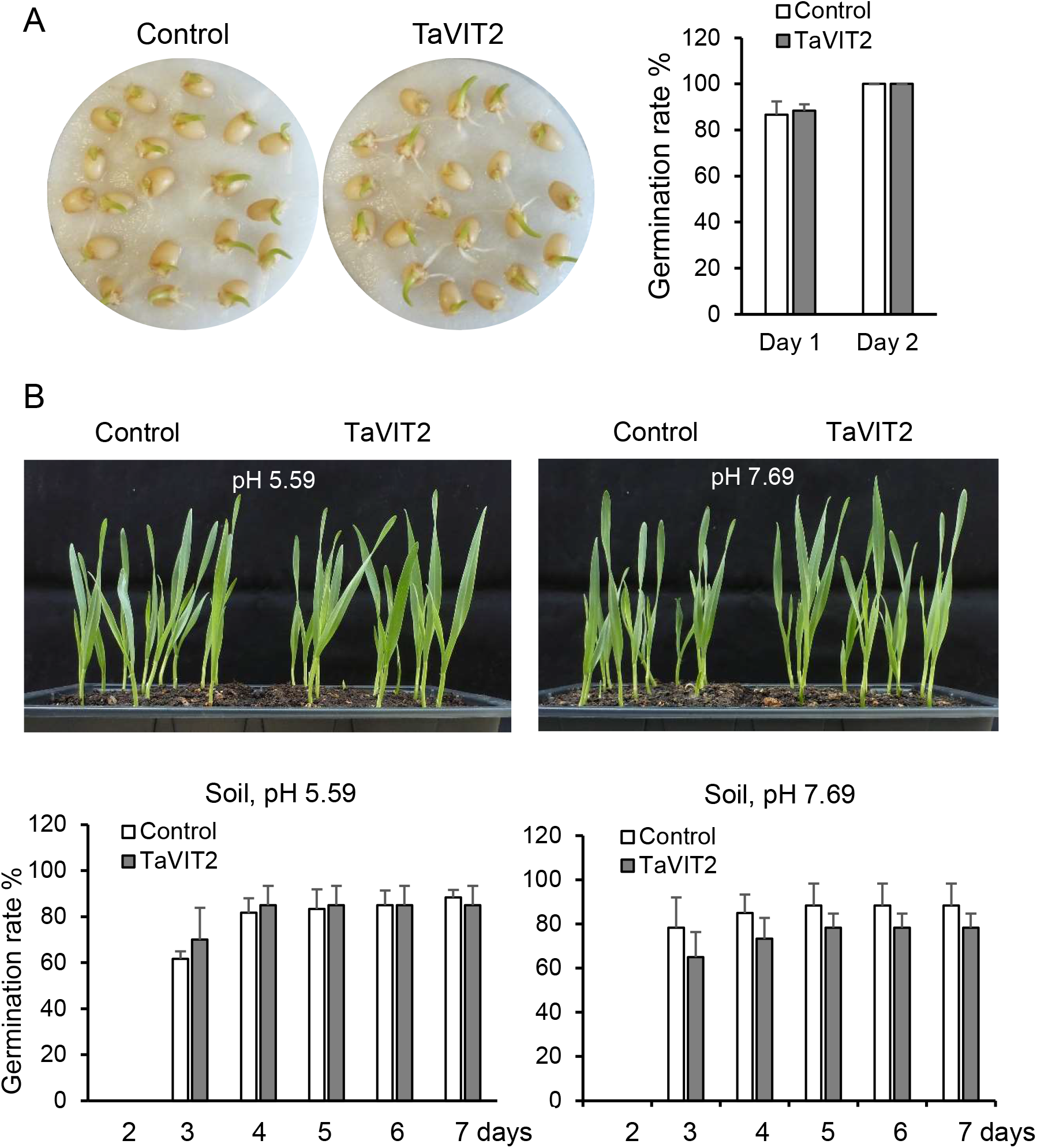
Germination of TaVIT2 grain is similar to control grain. **A**. Germination on cotton pads soaked with distilled water. Germination was scored as radical or shoot emergence >1 mm. The germination rate was calculated from 20 seeds in 3 replicates. **B**. Germination on normal and alkaline soil. Germination was scored as shoot emergence from 15 seeds in 4 replicates. For the graphs in (A) and (B), error bars represent SD. P > 0.1 for each pairwise comparison between control and TaVIT2 (non-significant, Student t-test).

**Supplemental Figure S3.**
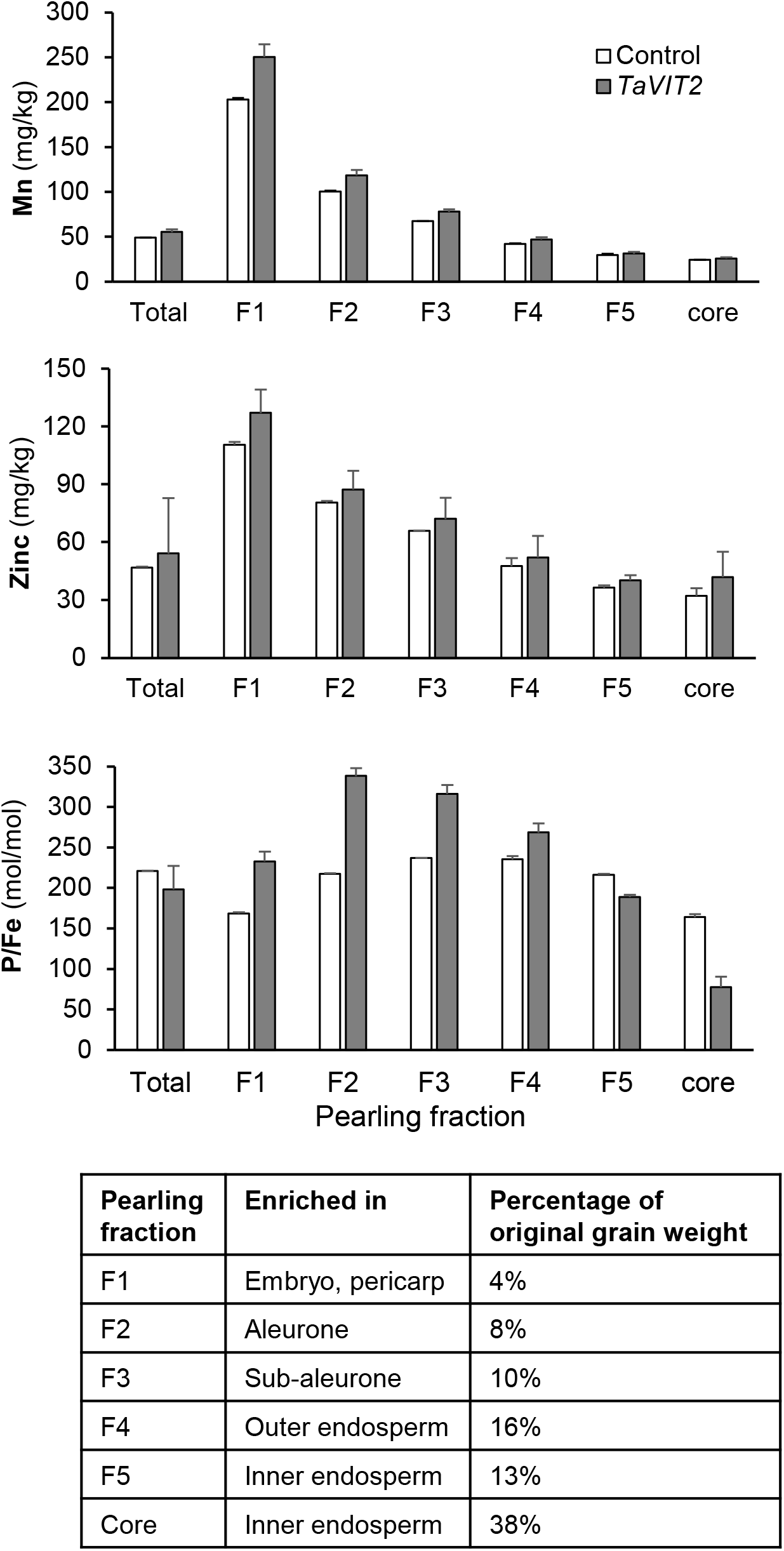
Element analysis in pearling fractions of wheat grain. Fractions obtained by pearling of mature grain were analysed by ICP-MS for manganese (Mn), zinc (Zn) and phosphorus (P). Values represent the mean of 2 biological replicates of 30 g grain. Error bars represent half the difference between the measurements. The tissue enrichment in each fractions is based on Tosi et al, (2018).

**Supplemental Figure S4.**
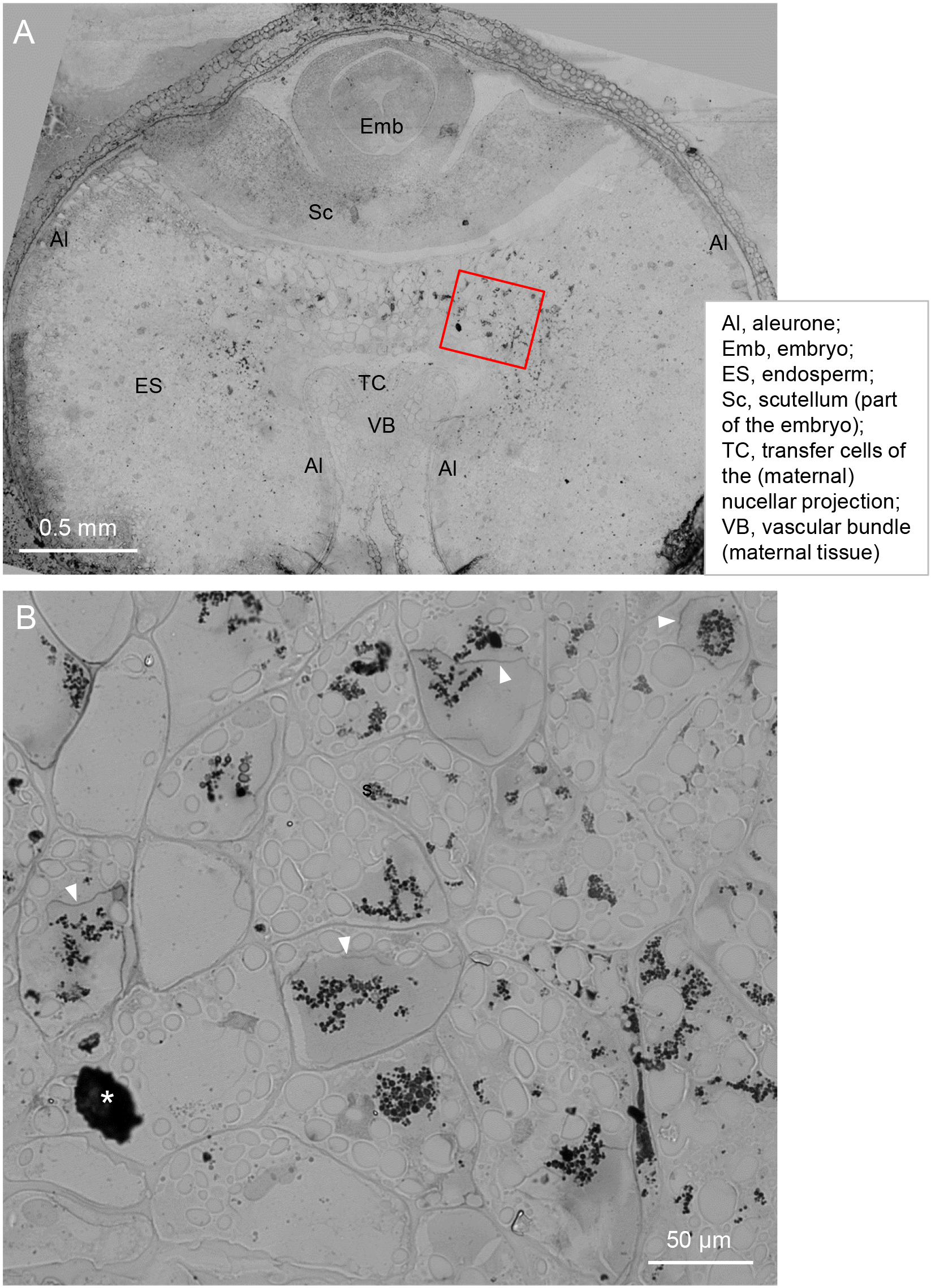
Additional images of enhanced Perls’ staining of TaVIT2 grain. **A**. Cross section through a grain harvested 21 dpa from wheat overexpressing *TaVIT2* from an endosperm-specific promoter. The section was stained for iron using the Perls’ method and enhanced with diamine benzidine. Images were taken with a 10x objectives lens and tiled. **B**. Enlarged image of the red rectangle in (A), taken with a 40x objective lens and DIC optics. (*) is a non-specific precipitate and should be ignored.

**Supplemental Figure S5.**
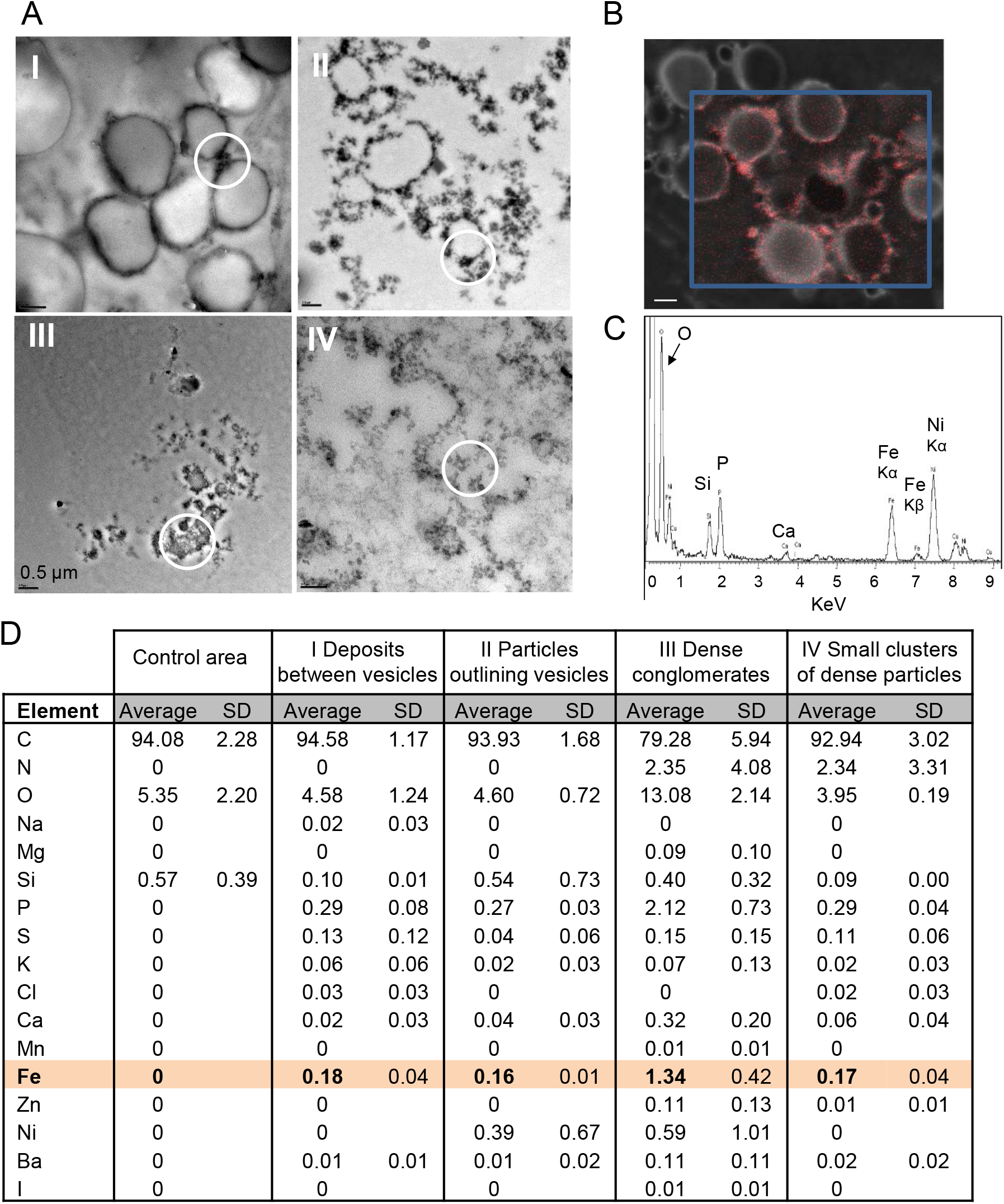
TEM and EDS analysis of TaVIT2 grain. **A**. Four different electron-dense morphologies (I – IV) were identified in endosperm cells adjacent to the scutellum (EAS) of TaVIT2 grain that were not found in control grain. The white circles in TEM images mark regions of interest (ROI) used for Energy Dispersive X-ray Spectroscopy (EDS) analysis (C and D). Scale bars are 0.5 μm. **B**. Elemental map obtained by STEM-EDS for morphology I, showing electron density and grey scale and iron in red. **C**. Example of an element profile obtained by EDS for morphology III, normalized for the Cu content of the TEM support grid. The peak at ~0.1 KeV, partially outside the y-axis scale, is carbon. **D**. Semi-quantitative analysis of EDS results. All values are percentage atomic weight, and the average of 3 areas for morphology I, II and IV, and 2 areas for III.

**Supplemental Figure S6.**
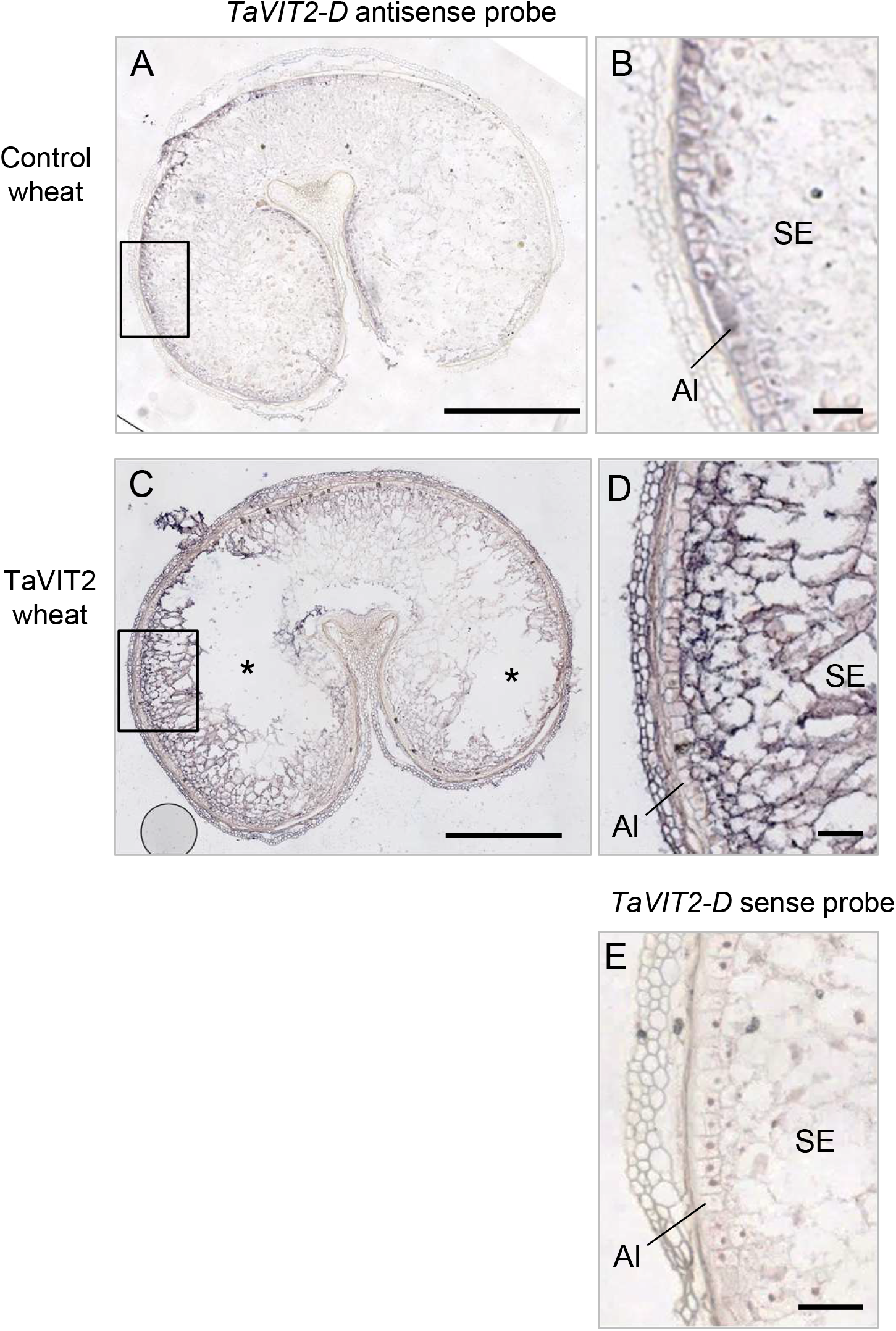
Expression pattern of the *HMW-TaVIT2* transgene. In-situ hybridization results for *TaVIT2-D* transcripts in cross sections of control wheat grains and from grains overexpressing TaVIT2-D under the control of the endosperm-specific *HMW Glu1-1D5* promoter. Grains were harvested at 21 dpa. **A**. Control grain hybridized with a DIG-labelled antisense RNA probe of TaVIT2-D. **B**. Larger magnification of the area indicated in (A). **C**. TaVIT2 grain hybridized with the same antisense RNA probe. White areas (*) is caused by tissue detachment from the slide. **D**. Larger magnification of the area indicated in (C). **E**. TaVIT2 grain hybridized with a sense RNA probe of *TaVIT2-D* as a negative control. Scale bars = 1 mm (A, C) or 0.1 mm (B, D and E). Al = aleurone, SE = starchy endosperm.

**Supplemental Figure S7.**
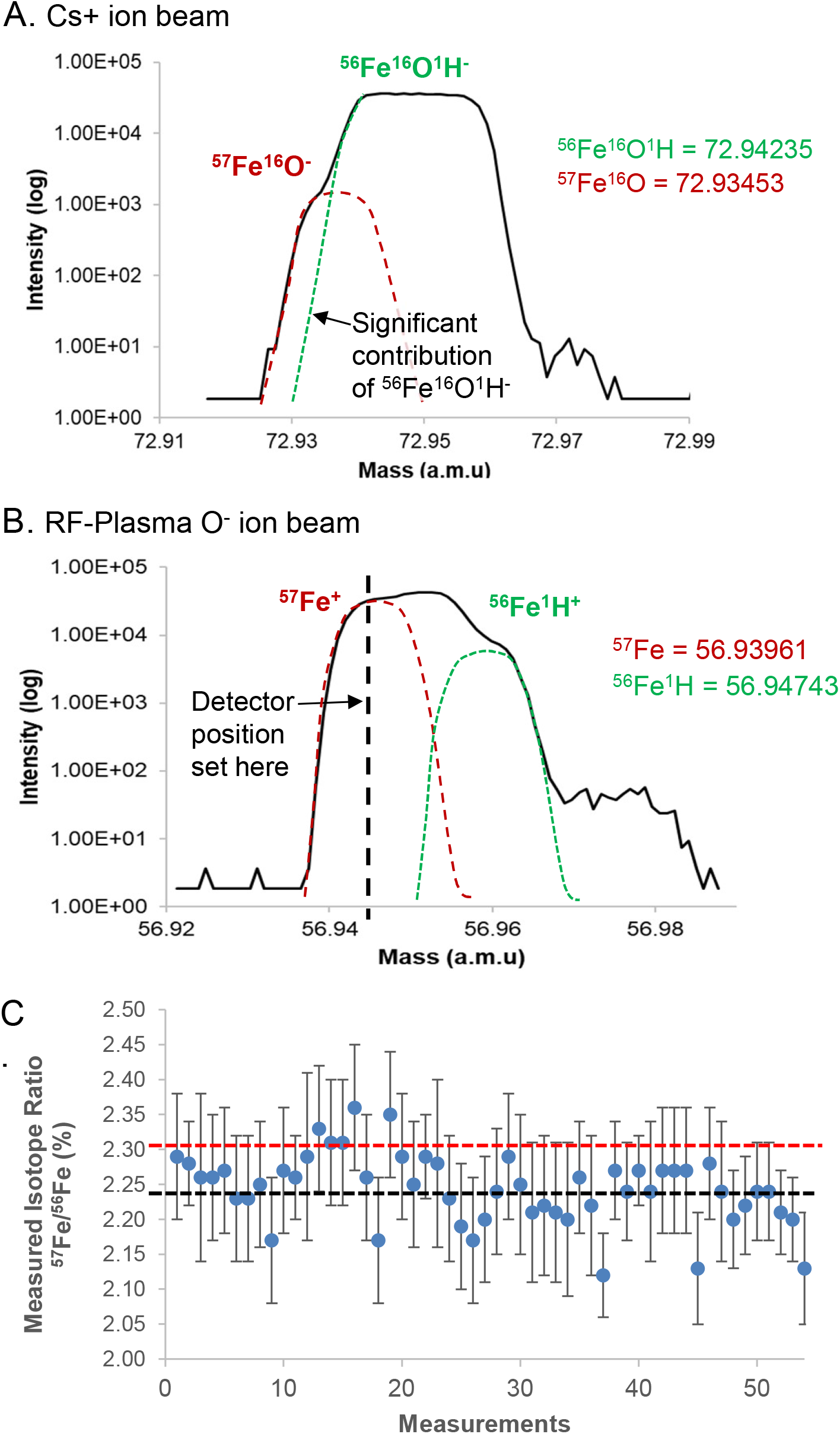
NanoSIMS data from the Fe standard. **A**. High-mass resolution scan using the Cs+ ion source, which shows the mass interference of 56Fe16O1H- and 57Fe16O-. **B**. High mass resolution scan using the RF-Plasma O-source, which shows that the detector can be positioned to separate 56Fe1H+ and 57Fe+ as indicated by the black dashed line. **C**. Ratio of 57Fe/56Fe measured on the Fe standard before each analysis. Orange line indicates the natural ratio at 2.31%; Black line indicates average ratio at 2.24%. Error bars show the Poisson error (%)

**Supplemental Figure S8.**
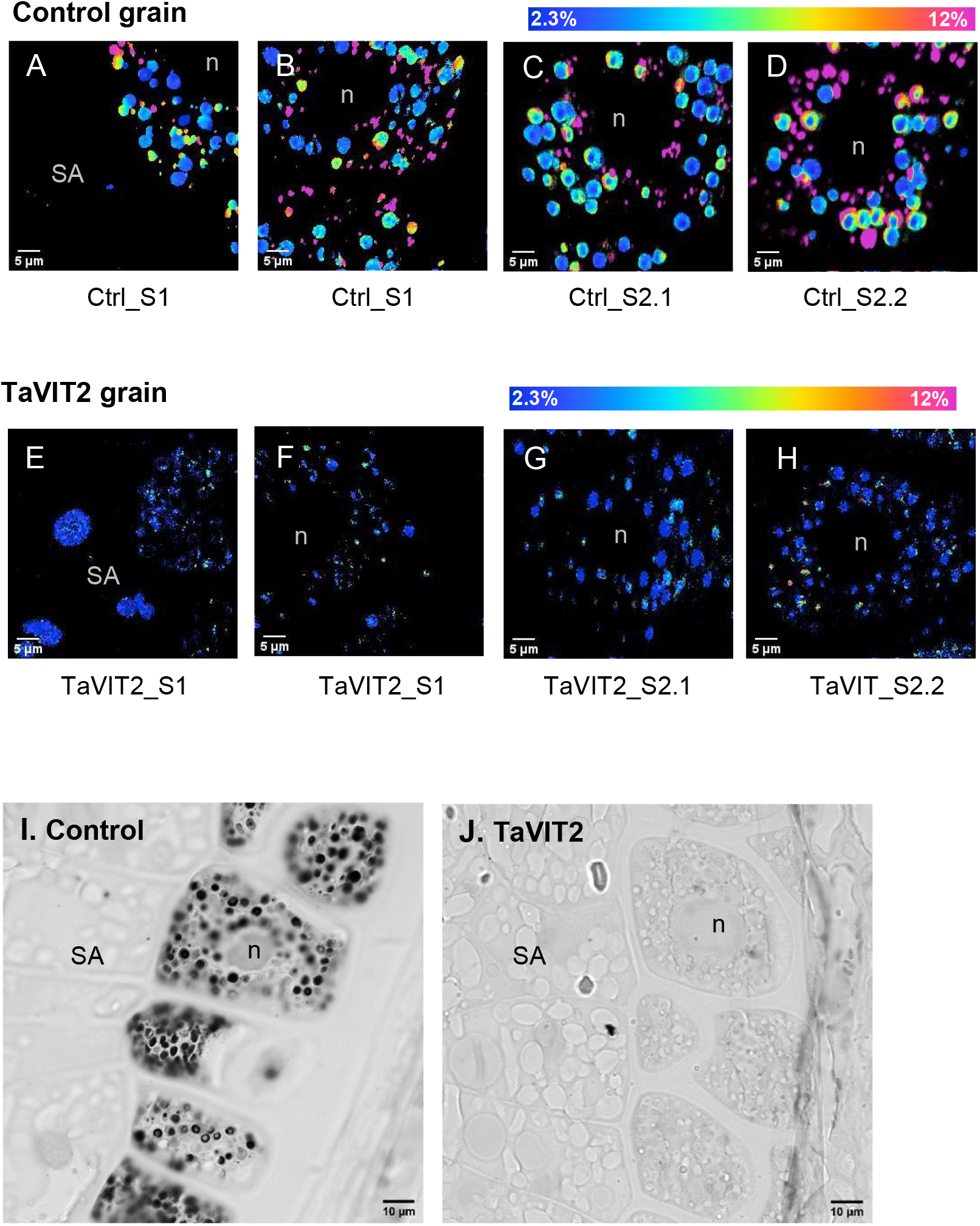
NanoSIMS images and chemical iron staining of aleurone cells. **A – H**. Overview of 50 × 50 μm NanoSIMS scans capturing (part of) aleurone cells. Control and TaVIT2 grains were labelled with 57Fe at 18 days post anthesis and analysed 72 hours later. The TaVIT2 line overexpresses a vacuolar iron transporter in the endosperm, limiting iron translocation to the aleurone cell layer. The images are from 2 biological replicates for each line: A and B are different areas of the same section; C and D are scans from two different sections of the same control grain. E, F, G and H are the analogous replicates in the TaVIT2 grain. SA, part of a subaleurone cell; n, nucleus. Scale bars for all images is 5 μm. I, J. Aleurone cells in control and TaVIT2 grain. Thin sections were stained for iron with enhanced Perls’ reagent and imaged by light microscopy.

**Supplemental Figure S9.**
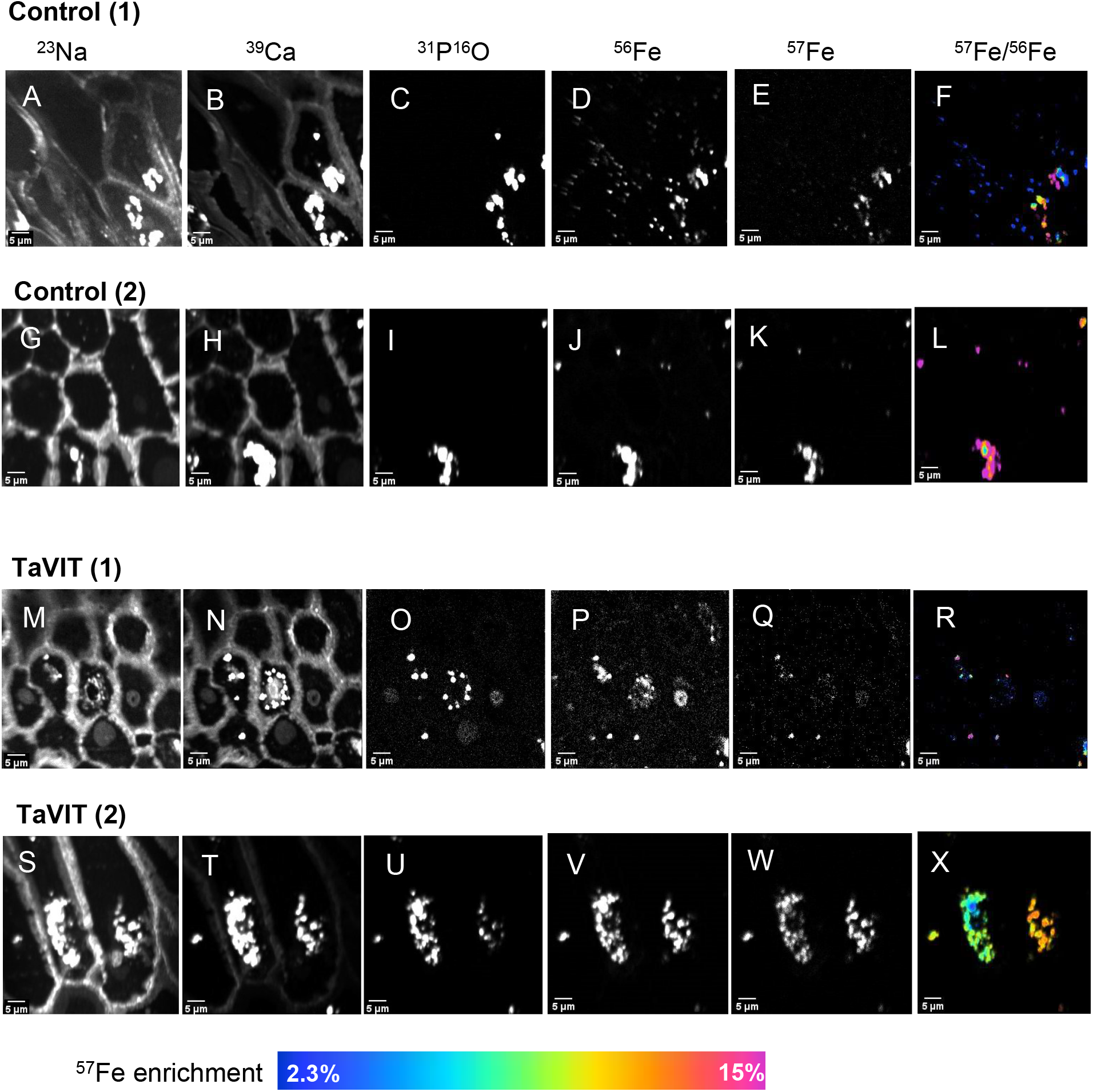
NanoSIMS images 50 × 50 μm of transfer cells of Control (A-L) and TaVIT2 (M-X) grain labelled with 57Fe at 18 days post anthesis and analysed 72 h later for the indicated ions. Images from 2 biological replicates are shown.

**Supplemental Figure S10.**
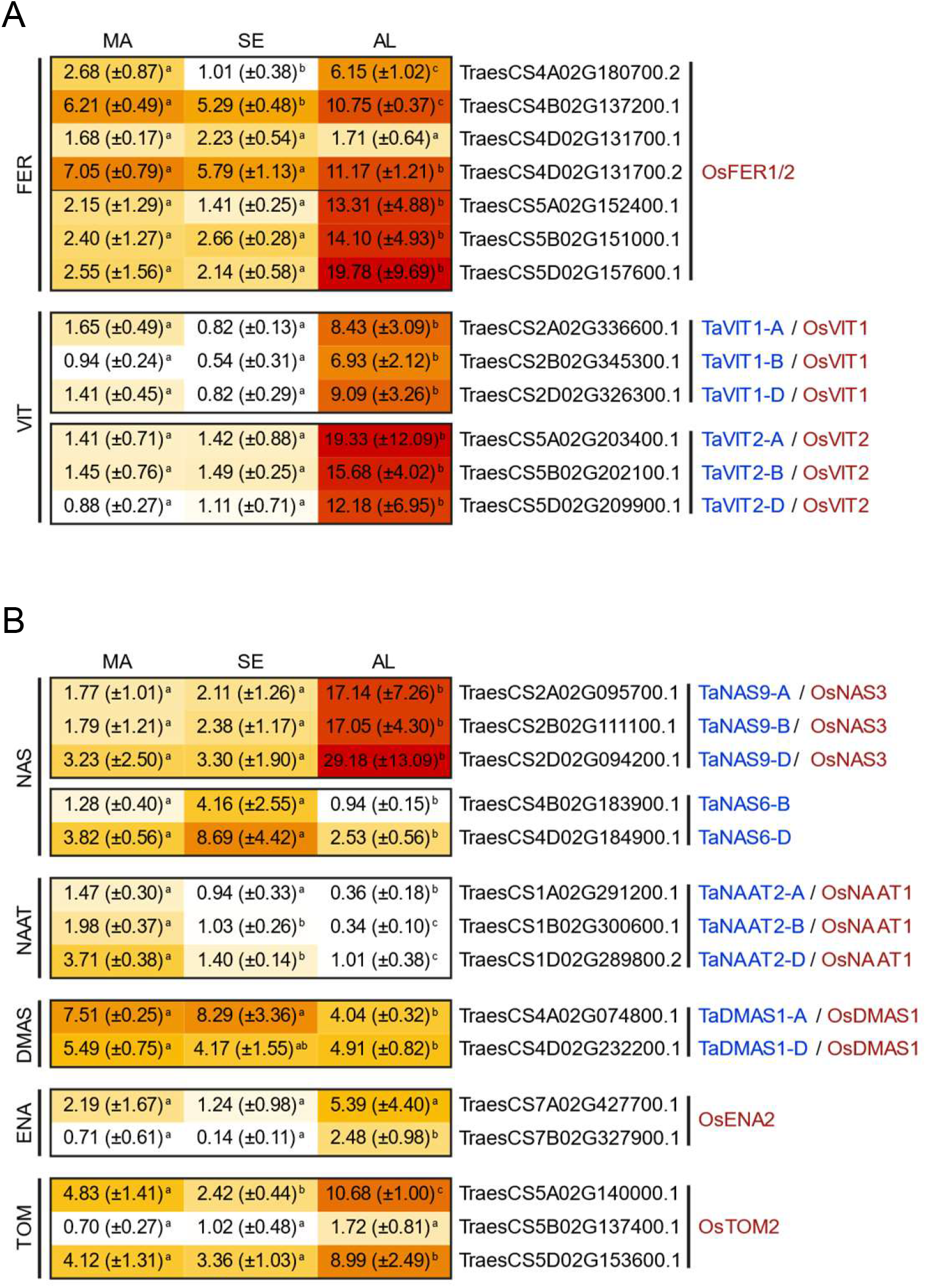
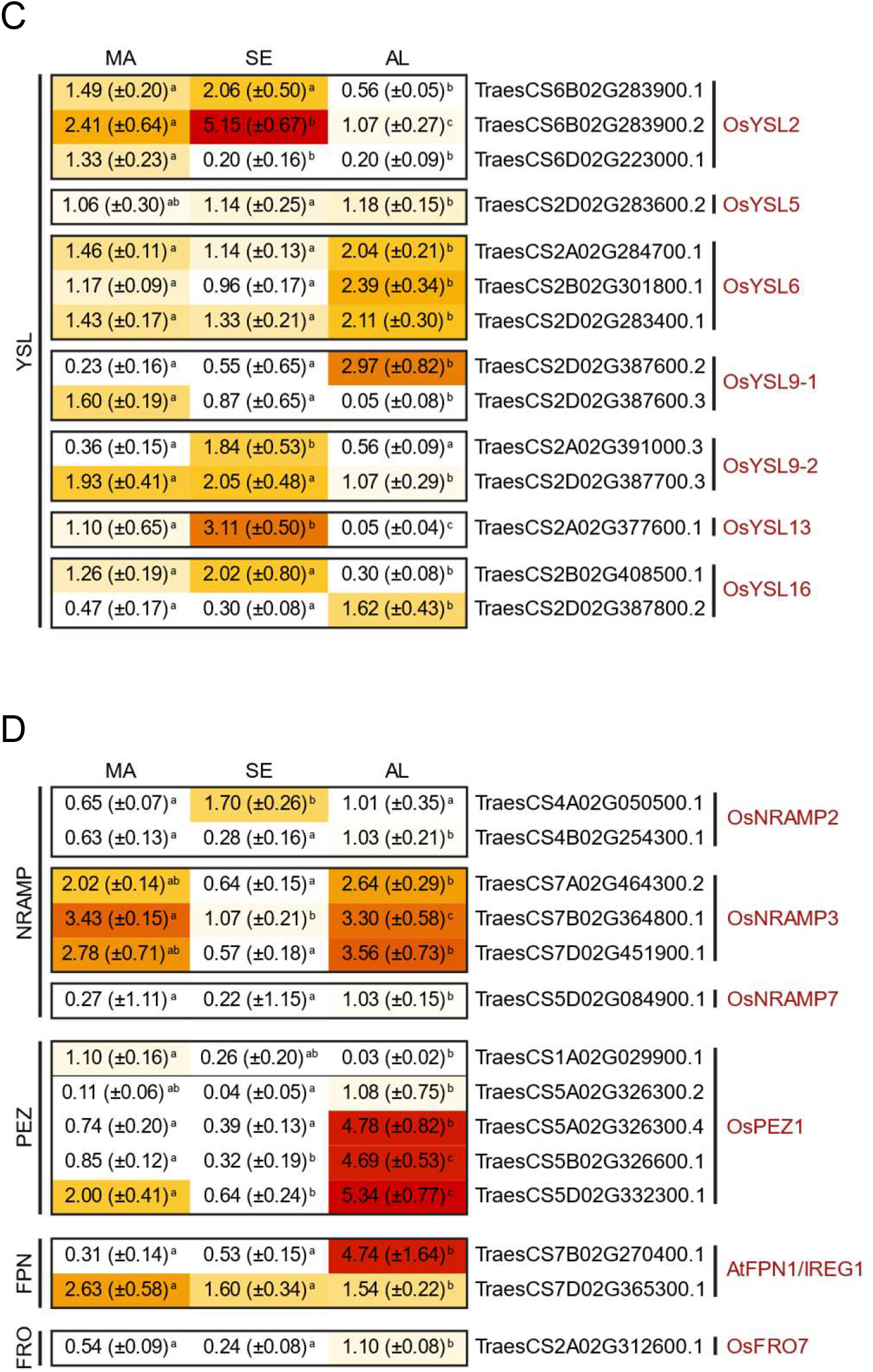
Transcript abundance of iron homeostasis genes in selected wheat grain tissues. RNA-seq data of samples enriched for modified aleurone cells (MA), starchy endosperm (SE) and aleurone cells (AL) of 20-dpa grain were from Pfeifer et al., 2014. Genes are grouped by their function: **A**. iron storage, **B**. nicotianamine and its derivatives biosynthesis and efflux, **C**. yellow stripe-like transporters, and **D**. other transporters putatively involved in iron homeostasis. Transcript identifiers are based on the International Wheat Genome Sequencing Consortium RefSeq v1.1. The values are transcripts per million (TPM) (± SD). Heatmap colouring is based on expression within each group (red showing highest expression within each panel).

**Supplemental Figure S11.**
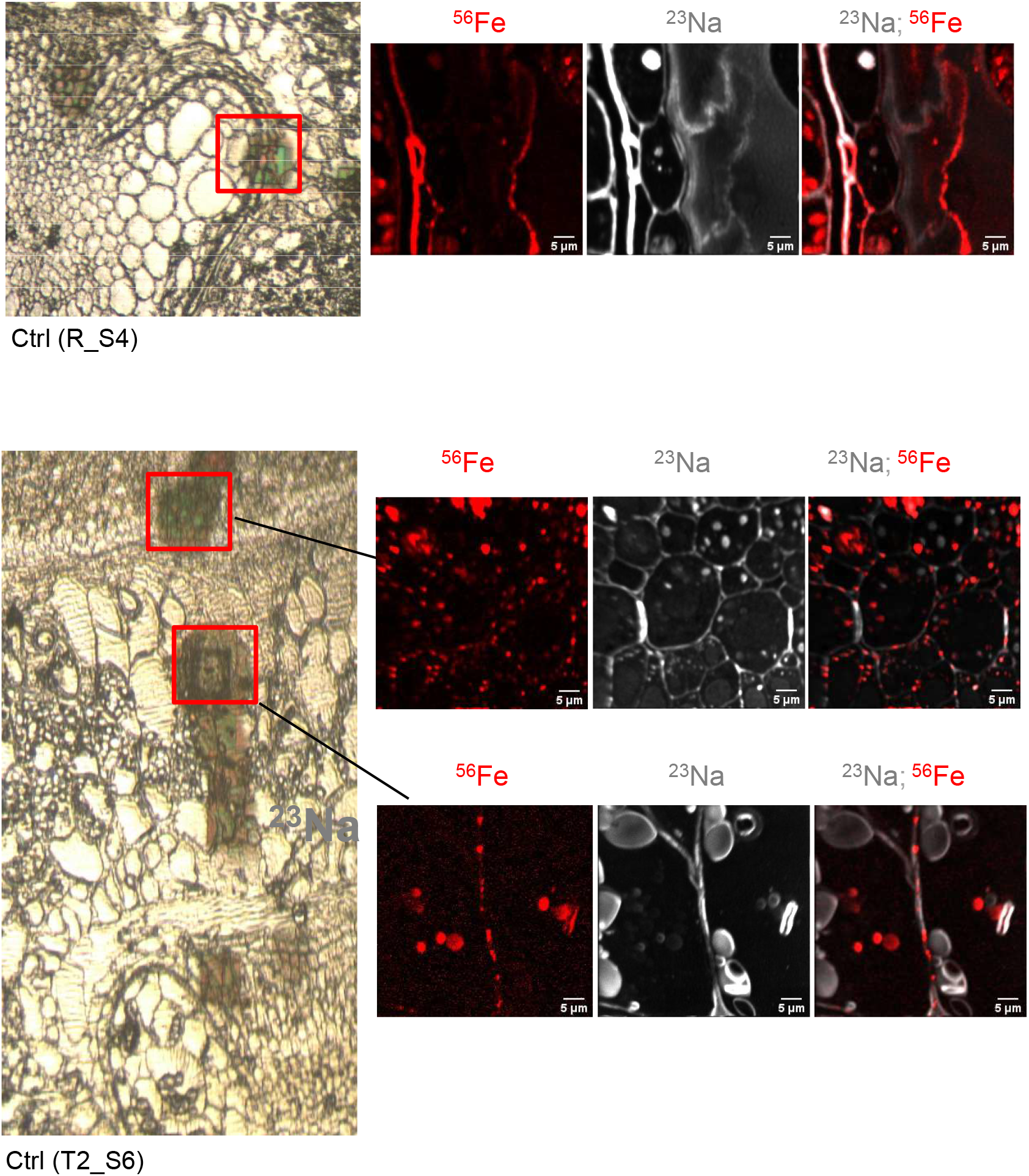
NanoSIMS images of iron in cell walls.

## Sheraz et al., Supplemental Methods

### Germination tests

Seeds were imbibed on damp filter paper and then grown on fine peat (90%) supplemented with 10% grit, 2.7 kg m^−3^ Osmocote, 3.25 kg m^−3^ Dolomitic limestone and 1 kg m^−3^ PG Mix (Yara, UK), which had a pH of ~5.6. To raise the pH for alkaline soil, 4.25 g kg^−1^ CaO was added. Plants were grown in a glasshouse kept at approximately 20°C with 16 h of light. Plants were watered as required.

### Element analysis by ICP-OES and ICP-MS

Preparation of a white flour fraction and subsequent inductively coupled plasma optical emission (ICP-OES) analysis were carried out as previously described (Connorton et al., 2017). In brief, grains were coarsely milled using a coffee grinder and then further ground using a mortar and pestle. The flour was passed through a 150-μm nylon mesh to remove most of the bran. Flour samples were dried overnight at 55°C and then digested overnight at 95°C in ultrapure nitric acid (55%, v/v) and hydrogen peroxide (6%, v/v). Samples were diluted 1:11 in ultrapure water and analysed by ICP-OES (Vista-PRO CCD Simultaneous ICP-OES; Agilent). As certified reference material we used Hard Red Spring Wheat Flour (REDS-1, National Research Council Canada) in parallel with all experimental samples.

Pearling of grain was carried out on 30 g grain samples in 2 replicates for each line using a Streckel and Schrader (Hamburg, Germany) pearling mill as described by Tosi et al. 2011. Five sequential cycles of pearling resulted in fractions F1−F5, which together accounted for about 50% of the grain weight. The remaining core is ~40% of the original weight, and ~10% is lost during milling. The grains remaining after pearling, called the core, were milled using a ball mill (Glen Creston, Stanmore, UK) and corresponded to about 50% of the original weight. Samples were dried at 80°C overnight and digested in ultrapure HNO_3_ and HClO_4_ (87:13% v/v) in triplicate using digestion blocks (200 °C, 12 h) (Eurotherm MBB151, Durrington, UK). Iron and other elements were measured by ICP-OES using a Perkin Elmer Optima 7300DV. The ^56^Fe and ^57^Fe iron isotopes in embryos and grain sectors were determined by Inductively Coupled Plasma – Mass Spectrometry (ICP-MS) using a Perkin Elmer NexION 300X.

*Scanning TEM imaging and EDS mapping* was done with a 200kV JEOL 2100PLUS TEM (JEOL Ltd, Tokyo, Japan) with a DF STEM detector and an Oxford Ultim Max EDS detector (Oxford instruments, Abingdon, UK), and results analysed with the Oxford Aztec software version 4.3.

### In-situ hybridization

Grains were harvested at 21 dpa, fixed in 4% (w/v) formaldehyde and embedded in paraffin wax as described in (Wan et al., 2014), except that Sub-X clearing agent (Leica) was used in place of HistoClear. Sections of 10 μm were hybridized with either an antisense or sense probe spanning nucleotides 1 – 738 of the *TaVIT2-5D* transcript (*TraesCS5D02G209900*). Hybridization was performed as described in (Wan et al., 2014), except that tissues were not acetylated prior to hybridization, and hybridization was carried out at 42°C instead of 50°C. Probes were labelled with digoxigenin (DIG) using the DIG RNA Labeling kit (Roche) and detected with anti-DIG antibody using the DIG Nucleic Acid Detection kit (Roche) according to manufacturer’s instructions.

### Sample preparation for nanoSIMS

Samples were transferred to tubes containing frozen 100% (v/v) ethanol in liquid nitrogen and placed in an automatic freeze substitution system (EM AFS from Leica Microsystems, UK). Samples were sequentially warmed over 5 days to −30 °C, then to 4 °C over 48 h and finally to room temperature. The samples were processed through increasing concentrations of LR White resin (Agar Scientific UK, R1281) and embedded at 58 °C for 16–20 h in a nitrogen-rich environment. Semi-thin sections (1 μm) of the resin blocks were cut with a Reichert-Jung ultramicrotome, and dried at 40 °C onto platinum-coated Thermanox coverslips for NanoSIMS. Adjacent sections were dried on Polysine Slides (Agar Scientific) for iron staining and for toluidine blue staining.

### Iron staining using the Perls’ method

Iron staining of tissue was performed as described in (Meguro et al., 2007; Roschzttardtz et al., 2009) using the Perls’ method. In brief, hand-cut wheat grains or 1-μm sections were incubated with 1:1 volumes of 4% (w/v) potassium ferrocyanide and 4% (w/v) HCl for 1 h and washed with water. For Perls’ stain intensification, samples were treated with methanol containing 0.01M NaN_3_ and 0.3% (v/v) H_2_O_2_ for 1h. After rinsing with 0.1 M phosphate buffer (pH 7.4) the sections were stained with 0.025% (w/v) 3,3’-diaminebenzidine 4-HCl, 0.005% (v/v) H_2_O_2_ and 0.005 (w/v) CoCl_2_ in 0.1 M phosphate buffer (pH 7.4) and washed with double distilled water.

## Sheraz et al., Supplemental Methods

### In-situ hybridization

Grains were harvested at 21 dpa, fixed in 4% (w/v) formaldehyde and embedded in paraffin wax as described in (Wan et al., 2014), except that Sub-X clearing agent (Leica) was used in place of HistoClear. Sections of 10 μm were hybridized with either an antisense or sense probe spanning nucleotides 1 – 738 of the TaVIT2-5D transcript (TraesCS5D02G209900). Hybridization was performed as described in (Wan et al., 2014), except that tissues were not acetylated prior to hybridization, and hybridization was carried out at 42°C instead of 50°C. Probes were labelled with digoxigenin (DIG) using the DIG RNA Labeling kit (Roche) and detected with anti-DIG antibody using the DIG Nucleic Acid Detection kit (Roche) according to manufacturer’s instructions.

## References

Bartlett, J., Alves, S., Smedley, M., Snape, J., and Harwood, W. (2008). High-throughput Agrobacterium-mediated barley transformation. Plant Methods 4: 22.

Bashir, K., Takahashi, R., Akhtar, S., Ishimaru, Y., Nakanishi, H., and Nishizawa, N.K. (2013). The knockdown of OsVIT2 and MIT affects iron localization in rice seed. Rice 6.

Beasley, J.T., Bonneau, J.P., and Johnson, A.A.T. (2017). Characterisation of the nicotianamine aminotransferase and deoxymugineic acid synthase genes essential to Strategy II iron uptake in bread wheat (Triticum aestivum L.). PLoS One 12: 1–18.

Beasley, J.T., Bonneau, J.P., Sánchez-Palacios, J.T., Moreno-Moyano, L.T., Callahan, D.L., Tako, E., Glahn, R.P., Lombi, E., and Johnson, A.A.T. (2019). Metabolic engineering of bread wheat improves grain iron concentration and bioavailability. Plant Biotechnol. J. 17: 1514–1526.

Bechtel, D.B., Abecassis, J., Shewry, P.R., and Evers, A.D. (2009). Development, structure, and mechanical properties of the wheat grain. In WHEAT: Chemistry and Technology, pp. 51–95.

Bonneau, J., Baumann, U., Beasley, J., Li, Y., and Johnson, A.A.T. (2016). Identification and molecular characterization of the nicotianamine synthase gene family in bread wheat. Plant Biotechnol. J. 14: 2228–2239.

Borg, S., Brinch-Pedersen, H., Tauris, B., and Holm, P.B. (2009). Iron transport, deposition and bioavailability in the wheat and barley grain. Plant Soil 325: 15–24.

Borrill, P., Connorton, J.M., Balk, J., Miller, A.J., Sanders, D., and Uauy, C. (2014). Biofortification of wheat grain with iron and zinc: Integrating novel genomic resources and knowledge from model crops. Front. Plant Sci. 5: 1–8.

Bray, N., Pimentel, H., Melsted, P., and Pachter, L. (2016). Near-optimal probabilistic RNA-seq quantification. Nat. Biotechnol. 34: 525–527.

De Brier, N., Gomand, S. V., Donner, E., Paterson, D., Smolders, E., Delcour, J.A., and Lombi, E. (2016). Element distribution and iron speciation in mature wheat grains (Triticum aestivum L.) using synchrotron X-ray fluorescence microscopy mapping and X-ray absorption near-edge structure (XANES) imaging. Plant Cell Environ. 39: 1835–1847.

Che, J., Yamaji, N., and Ma, J. (2021). Role of a vacuolar iron transporter OsVIT2 in the distribution of iron to rice grains. New Phytol.

Cominelli, E., Pilu, R., and Sparvoli, F. (2020). Phytic acid and mineral biofortification strategies: from plant science to breeding and biotechnological approaches. Plants (Basel) 9: 553.

Connorton, J.M., Jones, E.R., Rodríguez-Ramiro, I., Fairweather-Tait, S., Uauy, C., and Balk, J. (2017). Wheat vacuolar iron transporter TaVIT2 transports Fe and Mn and is effective for biofortification. Plant Physiol. 174: 2434–2444.

Detterbeck, A., Pongrac, P., Persson, D.P., Vogel-mikus, K., Kelemen, M., Husted, S., Schjoerring, J.K., Vavpetic, P., Pelicon, P., Arc, I., and Clemens, S. (2020). Temporal and spatial patterns of zinc and iron accumulation during barley (Hordeum vulgare L.) grain development. J. Agric. Food Chemsitry.

Doll, N.M., Just, J., Brunaud, V., Caïus, J., Grimault, A., Depège-Fargeix, N., Esteban, E., Pasha, A., Provart, N.J., Ingram, G.C., Rogowsky, P.M., and Widiez, T. (2020). Transcriptomics at maize embryo/endosperm interfaces identifies a transcriptionally distinct endosperm subdomain adjacent to the embryo scutellum. Plant Cell 32: 833–852.

Engelke, D.R., Krikos, A., Bruck, M.E., and Ginsburg, D. (1990). Purification of Thermus aquaticus DNA polymerase expressed in Escherichia coli. Anal. Biochem. 191: 396–400.

Evers, A.D. (1970). Development of the endosperm of wheat. Ann. Bot. 34: 547–555.

Garnett, T.P. and Graham, R.D. (2005). Distribution and remobilization of iron and copper in wheat. Ann. Bot. 95: 817–826.

Grovenor, C.R.M., Smart, K.E., Kilburn, M.R., Shore, B., Dilworth, J.R., Martin, B., Hawes, C., and Rickaby, R.E.M. (2006). Specimen preparation for NanoSIMS analysis of biological materials. Appl. Surf. Sci. 252: 6917–6924.

Hamdi, A., Roshan, T.M., Kahawita, T.M., Mason, A.B., Sheftel, A.D., and Ponka, P. (2016). Erythroid cell mitochondria receive endosomal iron by a “kiss-and-run” mechanism. Biochim. Biophys. Acta- Mol. Cell Res. 1863: 2859–2867.

Hayta, S., Smedley, M.A., Demir, S.U., Blundell, R., Hinchliffe, A., Atkinson, N., and Harwood, W.A. (2019). Correction to: An efficient and reproducible Agrobacterium- mediated transformation method for hexaploid wheat (Triticum aestivum L.) (Plant Methods (2019) 15: 121 DOI: 10.1186/s13007-019-0503-z). Plant Methods 15: 1–15.

Hoppe, P., Cohen, S., and Meibom, A. (2013). Technical aspects and applications in cosmochemistry and biological geochemistry. Geostand. Geoanalytical Res. 37: 111–154.

Ishimaru, Y., Kakei, Y., Shimo, H., Bashir, K., Sato, Y., Sato, Y., Uozumi, N., Nakanishi, H., and Nishizawa, N.K. (2011). A rice phenolic efflux transporter is essential for solubilizing precipitated apoplasmic iron in the plant stele. J. Biol. Chem. 286: 24649–24655.

Iwai, T., Takahashi, M., Oda, K., Terada, Y., and Yoshida, K.T. (2012). Dynamic changes in the distribution of minerals in relation to phytic acid accumulation during rice seed development. Plant Physiol. 160: 2007–2014.

Kim, S.A., Punshon, T., Lanzirotti, A., Li, L., Alonso, J.M., Ecker, J.R., Kaplan, J., and Guerinot, M.L. (2006). Localization of iron in Arabidopsis seed requires the vacuolar membrane transporter VIT1. Science (80-.). 314: 1295–1298.

Kopittke, P.M., Lombi, E., van der Ent, A., Wang, P., Laird, J.S., Moore, K.L., Persson, D.P., and Husted, S. (2020). Methods to visualize elements in plants. Plant Physiol. 182: 1869–1882.

Lamacchia, C., Shewry, P.R., Di Fonzo, N., Forsyth, J.L., Harris, N., Lazzeri, P.A., Napier, J.A., Halford, N.G., and Barcelo, P. (2001). Endosperm‐specific activity of a storage protein gene promoter in transgenic wheat seed. J. Exp. Bot. 52: 243–250.

Malherbe, J., Penen, F., Isaure, M.P., Frank, J., Hause, G., Dobritzsch, D., Gontier, E., Horréard, F., Hillion, F., and Schaumlöffel, D. (2016). A New Radio Frequency Plasma Oxygen Primary Ion Source on Nano Secondary Ion Mass Spectrometry for Improved Lateral Resolution and Detection of Electropositive Elements at Single Cell Level. Anal. Chem. 88: 7130–7136.

Mari, S., Bailly, C., and Thomine, S. (2020). Handing off iron to the next generation: How does it get into seeds and what for? Biochem. J. 477: 259–274.

Meguro, R., Asoano, Y., Odagiri, S., Li, C., Iwatsuki, H., and Shoumura, K. (2007). Nonheme-iron histochemistry for light and electron microscopy: a historical, theoretical and technical review. Arch. Histol. Cytol. 70: 1–19.

Moore, K.L., Tosi, P., Palmer, R., Hawkesford, M.J., Grovenor, C.R.M., and Shewry, P.R. (2016). The dynamics of protein body formation in developing wheat grain. Plant Biotechnol. J. 14: 1876–1882.

Moore, K.L., Zhao, F.-J., Gritsch, C.S., Tosi, P., Hawkesford, M.J., McGrath, S.P., Shewry, P.R., and Grovenor, C.R.M. (2012). Localisation of iron in wheat grain using high resolution secondary ion mass spectrometry. J. Cereal Sci. 55: 183–187.

Neal, A.L., Geraki, K., Borg, S., Quinn, P., Mosselmans, J.F., Brinch-Pedersen, H., and Shewry, P.R. (2013). Iron and zinc complexation in wild-type and ferritin-expressing wheat grain: Implications for mineral transport into developing grain. J. Biol. Inorg. Chem. 18: 557–570.

Olsen, O.A. (2020). The Modular Control of Cereal Endosperm Development. Trends Plant Sci. 25: 279–290.

Ondrasek, G., Rengel, Z., Clode, P.L., Kilburn, M.R., Guagliardo, P., and Romic, D. (2019). Zinc and cadmium mapping by NanoSIMS within the root apex after short-term exposure to metal contamination. Ecotoxicol. Environ. Saf. 171: 571–578.

Pfeifer, K., Kugler, K., Sandve, S., Zhan, B., Rudi, H., Hvidsten, T., IWGSC, Meyer, K., and Olsen, O. (2014). Genome interplay in the grain transcriptome of hexaploid bread wheat. Science (80-.). 345: 1250091.

Pongrac, P., Arcŏn, I., Castillo-Michel, H., and Vogel-mikuš, K. (2020). Mineral element composition in grain of awned and awnletted wheat (Triticum aestivum L.) cultivars: Tissue-specific iron speciation and phytate and non-phytate ligand ratio.

Pottier, M., Dumont, J., Masclaux-Daubresse, C., and Thomine, S. (2019). Autophagy is essential for optimal translocation of iron to seeds in Arabidopsis. J. Exp. Bot. 70: 845–858.

Raina, J.B. et al. (2017). Subcellular tracking reveals the location of dimethylsulfoniopropionate in microalgae and visualises its uptake by marine bacteria. Elife 6: e23008.

Roschzttardtz, H., Conéjéro, G., Curie, C., and Mari, S. (2009). Identification of the endodermal vacuole as the iron storage compartment in the Arabidopsis embryo. Plant Physiol. 151: 1329–1338.

Senoura, T., Sakashita, E., Kobayashi, T., Takahashi, M., Aung, M.S., Masuda, H., Nakanishi, H., and Nishizawa, N.K. (2017). The iron-chelate transporter OsYSL9 plays a role in iron distribution in developing rice grains. Plant Mol. Biol. 95: 375–387.

Singh, S.P., Vogel-Mikuš, K., Arčon, I., Vavpetič, P., Jeromel, L., Pelicon, P., Kumar, J., and Tuli, R. (2013). Pattern of iron distribution in maternal and filial tissues in wheat grains with contrasting levels of iron. J. Exp. Bot. 64: 3249–3260.

Takahashi, M., Nozoye, T., Kitajima, N., Fukuda, N., Hokura, A., Terada, Y., Nakai, I., Ishimaru, Y., Kobayashi, T., Nakanishi, H., and Nishizawa, N.K. (2009). In vivo analysis of metal distribution and expression of metal transporters in rice seed during germination process by microarray and X-ray Fluorescence Imaging of Fe, Zn, Mn, and Cu. Plant Soil 325: 39–51.

Uauy, C., Distelfeld, A., Fahima, T., Blechl, A., and Dubcovsky, J. (2006). A NAC gene regulating senescence improves grain protein, zinc, and iron content in wheat. Science (80-.). 314: 1298–1301.

Vasconcelos, M.W., Gruissem, W., and Bhullar, N.K. (2017). Iron biofortification in the 21st century: setting realistic targets, overcoming obstacles, and new strategies for healthy nutrition. Curr. Opin. Biotechnol. 44: 8–15.

Wang, Q., Zang, Y., Zhou, X., and Xiao, W. (2017). Characterization of four rice UEV1 genes required for Lys63-linked polyubiquitination and distinct functions. BMC Plant Biol. 17: 1–12.

Waters, B.M., Uauy, C., Dubcovsky, J., and Grusak, M.A. (2009). Wheat (Triticum aestivum) NAM proteins regulate the translocation of iron, zinc, and nitrogen compounds from vegetative tissues to grain. J. Exp. Bot. 60: 4263–4274.

WHO (2013). Research for universal health coverage.

WHO (2015). The global prevalence of anaemia in 2011.

Wu, L., Di, D.W., Zhang, D., Song, B., Luo, P., and Guo, G.Q. (2015). Frequent problems and their resolutions by using thermal asymmetric interlaced PCR (TAIL-PCR) to clone genes in Arabidopsis T-DNA tagged mutants. Biotechnol. Biotechnol. Equip. 29: 260–267.

Zang, J., Huo, Y., Liu, J., Zhang, H., Liu, J., and Chen, H. (2020). Maize YSL2 is required for iron distribution and development in kernels. J. Exp. Bot. 71: 5896–5910.

Zhang, Y., Xu, Y.H., Yi, H.Y., and Gong, J.M. (2012). Vacuolar membrane transporters OsVIT1 and OsVIT2 modulate iron translocation between flag leaves and seeds in rice. Plant J. 72: 400–410.

